# Acute psilocybin enhances cognitive flexibility in rats

**DOI:** 10.1101/2023.01.09.523291

**Authors:** Alejandro Torrado Pacheco, Randall J. Olson, Gabriela Garza, Bita Moghaddam

## Abstract

Psilocybin has been shown to improve symptoms of depression and anxiety when combined with psychotherapy or other clinician-guided interventions. To understand the neural basis for this pattern of clinical efficacy, experimental and conceptual approaches that are different than traditional laboratory models of anxiety and depression are needed. A potential novel mechanism is that acute psilocybin improves cognitive flexibility, which then enhances the impact of clinician-assisted interventions. Consistent with this idea, we find that acute psilocybin robustly improves cognitive flexibility in male and female rats using a task where animals switched between previously learned strategies in response to uncued changes in the environment. Psilocybin did not influence Pavlovian reversal learning, suggesting that its cognitive effects are selective to enhanced switching between previously learned behavioral strategies. The serotonin (5HT) 2A receptor antagonist ketanserin blocked psilocybin’s effect on set-shifting, while a 5HT2C-selective antagonist did not. Ketanserin alone also improved set-shifting performance, suggesting a complex relationship between psilocybin’s pharmacology and its impact on flexibility. Further, the psychedelic drug 2,5-Dimethoxy-4-iodoamphetamine (DOI) impaired cognitive flexibility in the same task, suggesting that this effect of psilocybin does not generalize to all other serotonergic psychedelics. We conclude that the acute impact of psilocybin on cognitive flexibility provides a useful behavioral model to investigate its neuronal effects relevant to its positive clinical outcome.

## Introduction

Recent clinical trials indicate that psilocybin is an effective treatment for symptoms of several psychiatric disorders, including anxiety and depression [1–5]. A unique aspect of these studies is that sustained therapeutic efficacy is observed when one or a few doses of psilocybin are administered during clinician-guided intervention [6–10]. To date, clinical data showing that psilocybin has therapeutic efficacy as a stand-alone treatment is lacking, suggesting that its effects may be, at least in part, due to the combination of therapeutic intervention and exposure to acute psilocybin. While the pharmacology of psilocybin is relatively well characterized, how this translates to its clinical effects remains unclear. What are the mechanisms that underlie psilocybin’s clinical efficacy when combined with psychotherapy or other clinician-guided interventions? Though several ideas have been proposed [11], we focused on the hypothesis that acutely psilocybin createsa cognitive state that improves therapeutic intervention by relaxing prior beliefs, promoting flexible thinking, and increasing cognitive flexibility [1,12,13].

Cognitive flexibility is the ability to switch between mental processes in order to appropriately adapt behavior to changes in the environment. Impairments in cognitive flexibility are a prominent feature of many psychiatric disorders, including major depression (MDD) and substance use disorders [14–18], which have shown a positive response to psilocybin-assisted treatment. Thus, the capacity to improve cognitive flexibility in order to facilitate breaking out of rigid behavioral patterns and beliefs is a plausible mechanism by which psilocybin may exert its therapeutic benefits [1,13,19].

Preclinical studies are critical for advancing our mechanistic understanding of the therapeutic efficacy of psilocybin. There is a growing appreciation that traditional behavioral tests used to study antidepressant effects do not reliably translate to effects in human populations [20]. While several traditional preclinical approaches to study depression-like behaviors in rodents have been used in conjunction with psilocybin [21,22], little is known about the acute effect of this drug on cognitive flexibility and related constructs. Other work has examined the acute effects of psychedelics on cognitive flexibility, but none of these studies used psilocybin [23,24]. We hypothesize that the acute effects of psilocybin may be inducing a state of increased cognitive flexibility that enhances the therapeutic response during clinician-assisted sessions.

To assess cognitive flexibility, we used a set-shifting task, which tests an animal’sability to modify behavior, among competing and previously learned strategies, in response to changes in environmental contingencies. We find that acute psilocybin treatment improves set-shifting behavior without influencing appetitive or aversive learning. Psilocybin’s effect on cognitive flexibility was diminished by a 5HT2A but not by a 5HT2C serotonin receptor antagonist. Consistent with other studies showing impaired cognitive flexibility with psychedelics other than psilocybin [23,24], we find that 2,5-Dimethoxy-4-iodoamphetamine (DOI), worsened set-shifting ability. These data support the idea that enhancing cognitive flexibility is a mechanism underlying the clinical effects of psilocybin.

## Materials and methods

### Animals

Experiments were performed in accordance with NIH’s *Guide to the Care and Use of Laboratory Animals* and were approved by the Oregon Health and Science University Institutional Animal Care and Use Committee. Long-Evans rats of both sexes were bred in house or purchased from Charles River Laboratories (CRL, Wilmington, MA) at 8 weeks of age. Rats (n=49, 24 males and 25 females) were pair-housed in same-sex pairs in a room with controlled humidity and temperature under a 12-h reverse light/dark cycle with lights off at 9:00 am. All behavioral tasks were conducted during the active dark phase. After acclimation and handling, animals were food restricted to 85-90% of their free-feeding weight before training on behavioral tasks began (see Supplementary Material for details of training procedures).

### Drugs

Psilocybin (received from Usona Institute and NIDA Investigational Drug and Material Supply Program) and DOI (Sigma-Aldrich, cat #D101) were dissolved in saline to a concentration of 1 mg/ml. Ketanserin (Sigma-Aldrich, cat #S006) and SB242084 (Tocris, cat #2901) were dissolved in 10% DMSO to a concentration of 1 mg/ml. Solutions were prepared on the day of injection when possible and otherwise stored at -20°C for up to a month. All injections were intraperitoneal. The dosage of 1 mg/kg for psilocybin and DOI has been shown to reliably elicit the head-twitch response, a behavioral marker of psychedelic activity [22,25]. The chosen dose of ketanserin abolishes the head-twitch responses produced by psilocybin [22]. We selected a relatively high dose of SB242084, which has behavioral effects in decision-making tasks [26,27]. Saline, DOI or psilocybin were injected 20 min before behavioral testing. Antagonist were injected 10 min before the main treatment (30 min prior to testing). Animals received multiple injections (min 1, max 4) given at least 2 weeks of washout between injections. Treatments were not repeated in the same animals and given in random order (Supplementary Table S1).

### Set-shifting task

Animals were tested on the extradimensional set-shifting task in daily sessions as previously described [28–30] (Fig. 1a; see Supplementary Materials for training procedure and testing details). Briefly, on each trial one of two nose-poke ports became illuminated. Rats were required to respond to one of the ports according to a “Light” rule, where the lit port was rewarded, or a “Side” rule, where the port on a specific side was rewarded, regardless of which one was lit. If and when rats reached a criterion of 10 consecutive correct responses, the rule changed. Each session lasted until criterion was reached on four rule sets, or until 45 minutes elapsed. Three separate cohorts of animals were used in this study, and behavior was consistent across cohorts (Supplementary Fig. S1).

**Figure 1.**
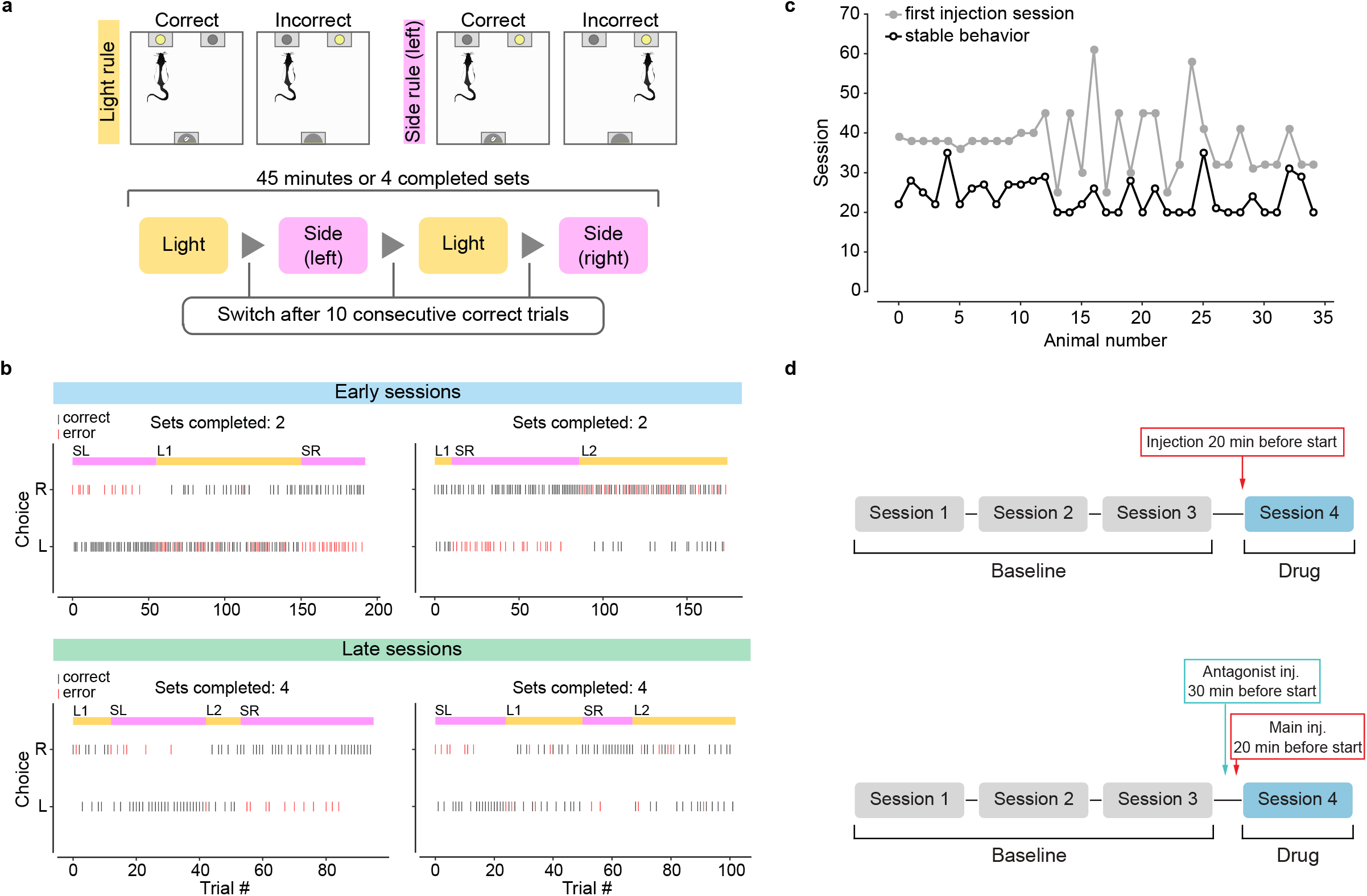
Set-shifting task. **(a)** Schematic representation of the operant set-shifting task. **(b)** Example sessions for one animal. Top, two sessions in the early stages of training. Bottom, two sessions during stable behavior. Vertical ticks denote the animal’s choice on each trial (Left or Right; black ticks, correct responses; red ticks, incorrect responses). Note that which port is illuminated on each trial is not shown. The current rule for each trial is shown in the colored bar at the top (yellow, Light rule; pink, Side rule). SL = side (left); SR = side (right); L1 = first light set; L2 = second light set. **(c)** Number of sessions until performance stabilized for every animal used in the set-shifting task (n=35, 18 males and 17 females). Black open circles indicate the point at which each animal reached stability (see Methods and Supplementary Materials for details). Gray solid circles indicate the first session in which that animal received a treatment injection (saline or drug). **(d)** Schematic of drug administration and task timeline. Three sessions of stable behavior were used as a baseline for each animal. For main treatment sessions (top) injections were given 20 minutes prior to the start of the session. For antagonist sessions (bottom), the antagonist drug was injected 10 minutes prior to the main treatment.

### Reversal learning task

The reversal learning task was a modified version of a previously characterized task [31] (see Supplementary Materials for more details). Briefly, two different conditioned stimuli were associated with an appetitive or aversive unconditioned stimulus (US). Animals performed ten task sessions during which 50 stimulus-outcome associations for each kind of CS-US pairing were presented. On the sixth session, CS-US contingencies were reversed such that the CS that previously signaled an appetitive outcome now predicted the aversive one, and vice-versa. Injections of psilocybin (1 mg/kg) or an equivalent volume of saline were given 20 min before starting the task on the reversal session (session 6). We chose to give only one dose of psilocybin for consistency with the set-shifting experiment.

### Analysis and statistical tests

To assess set-shifting performance we calculated trials to criterion, number of completed sets, and fraction of correct responses, for 3 days of baseline and for the drug treatment session (Fig. 1d). The average trials to criterion value was obtained by calculating the number of trials needed to reach 10 consecutive correct responses on each set and averaging across all completed sets in a session. Note that this excludes sets that were not completed and therefore may be artificially low when animals quickly complete one set but then fail to shift their behavior to the new rule. This would result in a low value for the average trials to criterion, but it cannot be taken as evidence of effective set-shifting behavior. For this reason, we used a combination of trials to criterion, number of completed sets, and fraction of correct responses to assess set-shifting performance. The fraction of correct responses was calculated by combining trials from all sets. We also calculated “streaks”, defined simply as any consecutive bout of correct responding. Lastly, we computed response time, defined as the time elapsed between the stimulus onset (lit nose-poke port) and action choice (nose-poke action). For the reversal learning task, we evaluated the behavior by computing the R-value [31]. This is defined for each stimulus in a session as the total number of responses in the food port during the cue presentations, divided by total number of responses in the food port during the second half of the ITIs. Because for each CS there are twice as many ITIs as there are CS presentations, a R-value of 0.5 corresponds to no conditioning. Larger values indicate appetitive conditioning, smaller values indicate aversive conditioning.

Statistical analyses were performed using GraphPad Prism software (GraphPad, San Diego, CA) or custom Python scripts. For all statistical tests we used t-tests when data were normally distributed and rank-sum or sign-rank tests when they were not. Two-way ANOVA (or mixed-effects analysis, when some data were missing) followed by Sidak or Dunnett’s post-hoc tests was used to compare the effect of two factors on behavior. For the reversal learning task, we used a mixed-effects model followed by Sidak post-hoc tests. The significance threshold for all tests was set to 0.05.

## Results

### Characterization of behavior in extradimensional set-shifting task

The effect of psilocybin on cognitive flexibility was tested using a previously characterized operant set-shifting task [28,30]. Adult rats of both sexes (n=35, 18 males and 17 females) were trained in the task and all animals learned the task well (Fig. 1; see Methods and Supplemental Methods for details). After animals reached stable behavior (Fig. 1c; see Supplementary Methods), they received their first i.p. injection 20 min before behavioral testing (Fig. 1d; see Supplementary Methods).

### Acute psilocybin improves performance in set-shifting task

Psilocybin (1 mg/kg, i.p.) improved performance on the set-shifting task, as indicated by a significant decrease in the average number of trials rats needed to reach criterion (Fig. 2a, p=0.005, n=15; see also Supplementary Fig. S2a, b). A comparable number of sets was completed with psilocybin treatment and baseline (Fig. 2b, p=0.546), despite significantly slower reaction times (Fig. 2c, p=0.008). Psilocybin did not change the fraction of correct responses (Fig. 2d, p=0.188) but correct responses can be made without reaching the criterion of 10 consecutive ones. We, therefore, analyzed the effect of psilocybin on a “streak”, defined as any series of consecutive correct responses. We observed that animals receiving psilocybin performed longer without making an incorrect response, resulting in fewer and longer streaks on average (Fig. 2e, f; number of streaks, p=0.025; average streak length, p=0.041). These analyses suggest that psilocybin selectively reduces incorrect responses occurring after a series of correct ones, thus increasing the likelihood of reaching criterion on the task.

**Figure 2.**
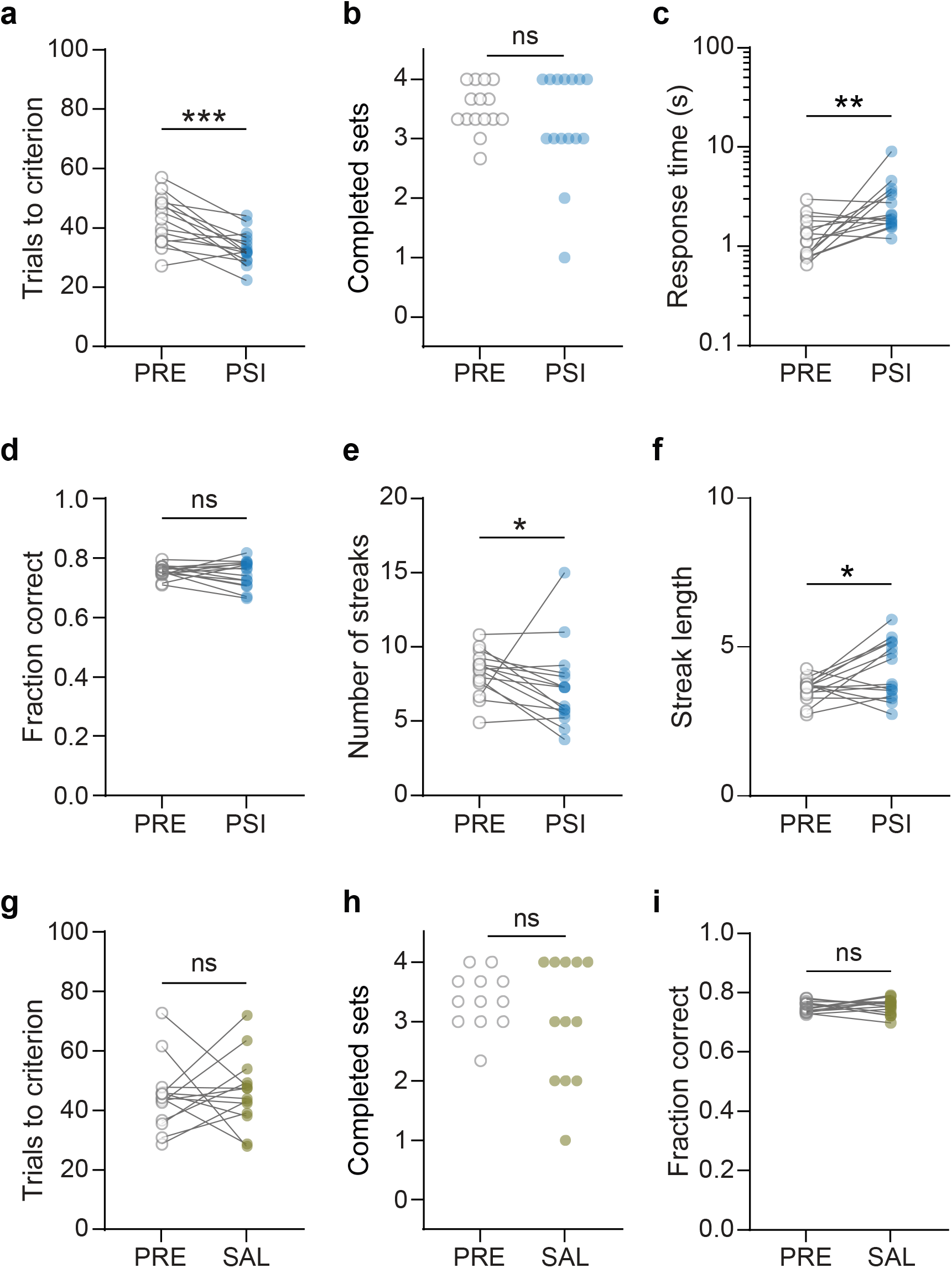
Psilocybin improves performance in the set-shifting task. **(a)** Average trials-to-criterion across all sets in the baseline period (PRE, open gray circles) and in the acute psilocybin condition (PSI, blue circles, n=15). *** p=0.005, paired t-test. **(b)** Number of completed sets in baseline and psilocybin conditions. **(c)** Response time in baseline and psilocybin conditions. ** p=0.008, Wilcoxon sign-rank test. **(d)** Fraction of correct responses across all sets in baseline and psilocybin conditions. **(e)** Number of streaks of consecutive correct responses averaged across all sets, in baseline and psilocybin conditions. * p=0.025, Wilcoxon sign-rank test. **(f)** Average length of streaks averaged across all sets, in baseline and psilocybin conditions. * p =0.034, paired t-test. **(g)** Average trials-to-criterion for baseline (PRE, gray open circles) and saline conditions (SAL, brown circles, n=14). **(h)** Number of completed sets in baseline and saline conditions. **(i)** Fraction of correct responses across all sets in baseline and saline conditions.

We performed a number of additional analyses to investigate the effects of psilocybin in detail. First, we note that the effect of psilocybin on trials to criterion was most pronounced in the first two sets (Supplementary Fig. S2a), possibly due to it wearing off in sets 3 and 4. To confirm that animals completed the first two sets while on the acute effects of psilocybin, we computed the time to criterion for each set during baseline and psilocybin days. Time to criterion values were similar in baseline and psilocybin days. Furthermore, nearly all animals completed the first two sets within 30 min of starting the task, i.e. within 50 min of the injection (Supplementary Fig. S2e), which is well within the window for psilocybin’s acute effects. Secondly, we analyzed Light- and Side-rule sets separately to ask whether psilocybin differentially affected performance depending on rule, rather than order of sets. These comparisons did not reach significance when looking at all four sets (data not shown; Light, p=0.059; Side, p=0.051), but they were both significant when we restricted the analysis to the first two sets (Supplementary Fig. S2f; Light, p=0.026; Side, p=0.024). We conclude that psilocybin had a similar effect on both Light and Side sets. Finally, we note that data for both sexes were combined as no sex differences were found (Supplementary Fig. S2g).

In the analyses above, we compared behavior on the day of treatment to the average performance in the previous three days (baseline days). It is critical for this analysis that the behavior in the baseline period is stable. To ensure that our stability criteria (see Supplementary Methods) resulted in stable baseline performance, we plotted trials to criterion, completed sets and fraction of correct responses on baseline, and on treatment days, for all psilocybin-treated animals. The behavior of individual animals oscillated within a consistent range across the baseline sessions, and the average across the cohort was stable (Supplementary Fig. S2b). We also aimed to control for rule preference. For each animal we split sessions by whether their starting rule (i.e. in the first set) was a light or side rule, and quantified behavioral metrics for stable sessions performed by that animal. This analysis showed that for the psilocybin-treated cohort, performance as a group was not biased by the starting rule in each session (Supplementary Fig. S2b).

Animals receiving vehicle saline injections completed a similar number of sets as they did during the baseline period (Fig. 2h, p=0.237, n=14; see also Supplementary Fig. S2c, d), but they did not show any difference in the trials-to-criterion metric on the day of injection (Fig. 2g, p=0.703). They also showed no change in the fraction of correct responses (Fig. 2i, p=0.463). We used the saline-treated group as a control and compared trials to criterion during baseline for all treatment groups to the baseline value for the saline group. We found no difference, indicating that all treatment groups had similar pre-treatment performance in the task (Supplementary Fig. S2h; two-way ANOVA followed by Dunnett’s test, all p>0.35). We also compared each group’s treatment data with that of the saline group and found a significant difference for psilocybin (Supplementary Fig. S2h; two-way ANOVA followed by Dunnett’s test, p=0.005), among others (see below).

### Psilocybin does not impact performance on a reversal learning task

Next we asked whether psilocybin can facilitate learning of changes in environmental contingencies, a construct generally assessed in reversal learning tasks. Reversal learning and set-shifting involve different neural mechanisms, consistent with the substantial difference in the cognitive demands of each task [32,33]: whereas reversal learning involves updating cue-outcome (or action-outcome) associations, set-shifting requires no new learning but rather tests the capacity to adaptively switch between known behavioral strategies. To test reversal learning, we used another previously characterized task [31]. This task has the advantage of assessing reversal learning related to both appetitive and aversive outcomes so that the impact of psilocybin on affective context could also be evaluated. Drug-naïve and untrained rats were exposed to two different cue-stimulus pairings over 10 task sessions. The cue was either a tone or a flashing light, and each was paired either with a mild foot-shock or a food reward (Fig. 3a; see Supplementary Methods for details). The pairings were switched on session 6. Rats were injected with psilocybin (1 mg/kg, n=7, 3 males and 4 females) or an equivalent volume of saline (n=6, 3 of each sex) 20 min before the reversal session (Fig. 3b).

**Figure 3.**
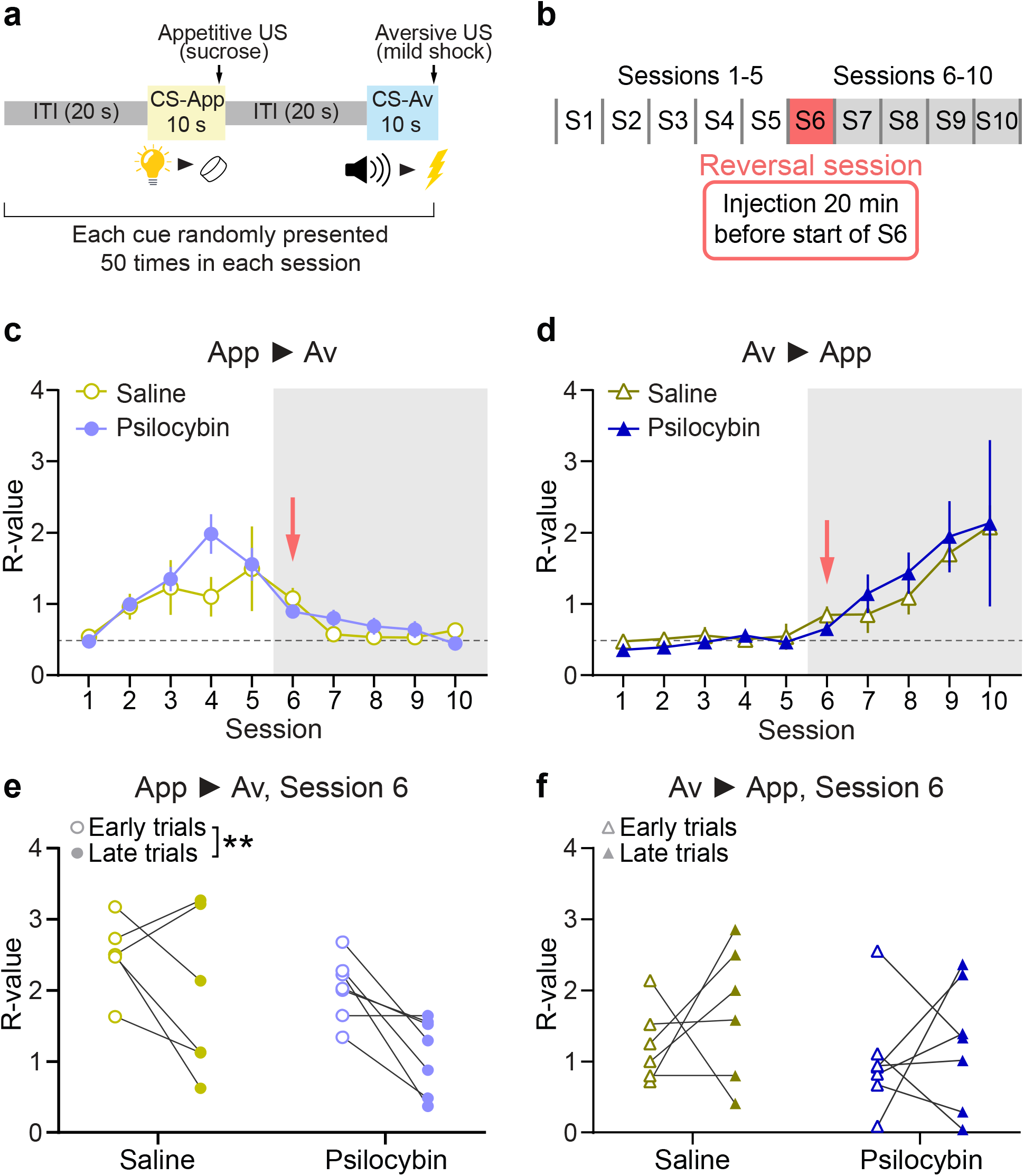
No effect of psilocybin in the reversal learning task. **(a)** Schematic diagram of the Pavlovian reversal learning task. **(b)** Structure of the reversal learning task, including time of injection. **(c)** R-value comparison of responding to the initially appetitive cue (App > Av) between saline-treated (open circles, n=6) and psilocybin-treated (solid circles, n=7) animals. Red arrow indicates the session in which contingencies change and treatment is administered. Mixed-effects analysis: main effect of Treatment, p=0.472; interaction effect, Session x Treatment, p=0.179. **(d)** As in (c), for the initially aversive cue. Mixed-effects analysis: main effect of Treatment, p=0.834; interaction effect, Session x Treatment, p=0.993. **(e)** R-value in early (first 10) and late (last 10) trials in session 6 of the reversal task, when the reversal happened, for the initially appetitive cue. All individual animals are shown. Two-way ANOVA: main effect of Timepoint, p=0.009; interaction effect, Treatment x Timepoint, p=0.487. **(f)** As in (e), for the initially aversive cue. Two-way ANOVA: main effect of Timepoint, p=0.371; interaction effect, Treatment x Timepoint, p=0.754.

Both saline and psilocybin groups learned the cue-outcome contingencies, and after the reversal the previously aversive cue became appetitive for both groups, and vice versa for the initially appetitive cue (Fig. 3c, d). No differences were found when comparing responses to each contingency pairing for the two treatment groups (Fig. 3c, d). To investigate differences in within-session learning during the reversal session, we plotted the R-value in early (first 10) and late (last 10) trials in that session, for each animal, cue type and treatment group (Fig. 3e, f). Both saline- and psilocybin-treated animals showed evidence of learning for the previously appetitive cue (Fig. 3e; two-way ANOVA, main effect of Timepoint, p=0.009), but there were no differences between groups (interaction effect Treatment x Timepoint, p=0.487). We also found no differences in within-session learning between groups for the previously aversive cue (Fig. 3f; two-way ANOVA, interaction effect Treatment x Timepoint, p=0.754).

These data show that psilocybin does not influence this form of reversal learning, regardless of the nature of the outcome, indicating specificity in its cognitive effects. Our results suggest that while acute psilocybin improves switching between known behavioral strategies, it does not improve learning of changes in environmental contingencies.

### Enhanced cognitive flexibility by psilocybin does not generalize to the psychedelic DOI and is blocked by ketanserin

To test whether enhanced flexibility could result from treatment with a different psychedelic agent, we treated animals with DOI, a commonly used psychedelic compound which is an agonist at 5HT2A receptors. DOI (1 mg/kg, i.p.) did not improve set-shifting performance (Fig. 4a, p=0.112, n=12; see also Supplementary Fig. S3). Moreover, rats completed significantly fewer sets when treated with DOI compared to the baseline period (Fig. 4b, p=0.012), with most completing only one or no sets at all. The fraction of correct responses was also lower compared to baseline (Fig. 4c, p=0.005). In the comparison to saline, the DOI group showed a significant effect (Fig. S2h, p=0.013). However, we argue that the low value for trials to criterion in the DOI group is largely due to artificially low numbers for that metric when animals complete very few sets, as explained above. Most animals treated with DOI only completed one set or none at all. Overall this represents a clear deficit in performance of extradimensional shifts. Thus, the effect of psilocybin on set-shifting does not generalize to all other serotonergic psychedelics, suggesting that psilocybin may have a unique profile of cognitive effects.

**Figure 4.**
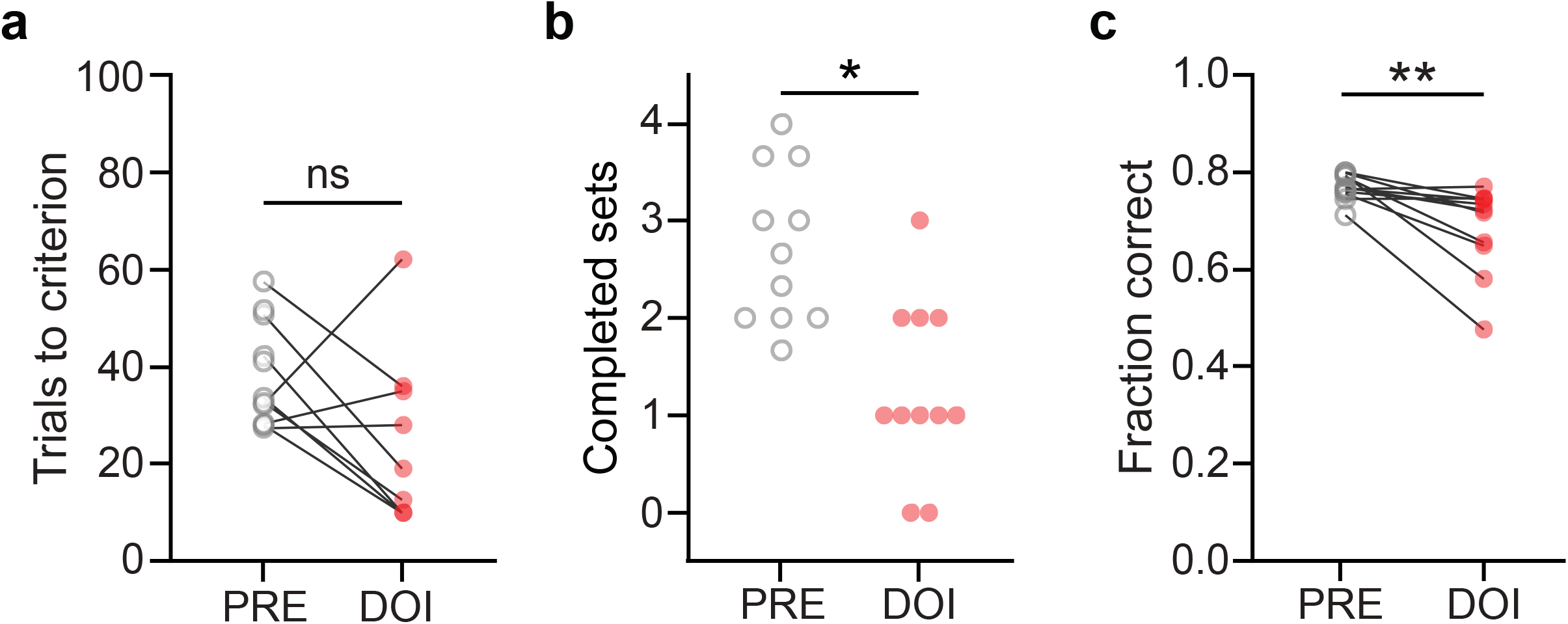
Treatment with DOI impairs performance in the set-shifting task. **(a)** Average trials-to-criterion for baseline (PRE, gray open circles) and DOI conditions (red circles, n=12). **(b)** Number of completed sets in baseline and DOI conditions. * p=0.012, Wilcoxon sign-rank test. **(c)** Fraction of correct responses averaged across all sets in baseline and DOI conditions. ** p=0.005, Wilcoxon sign-rank test.

Psilocin, the active metabolite of psilocybin, is a partial agonist of both 5HT2A and 5HT2C receptors. Activation of the former is thought to be responsible for the subjective effects of psilocybin and other psychedelics, so we investigated whether agonism of 5HT2A receptors played a role in the cognitive effects we observed. We pre-treated ratswith the 5HT2A antagonist ketanserin (1 mg/kg, i.p.) 10 min before injection of either psilocybin or saline. Ketanserin blocked the effect of psilocybin on cognitive flexibility (Fig. 5a-c, n=14; trials to criterion, p=0.231; completed sets, p=0.688; fraction correct, p=0.469; see also Supplementary Fig. S4). This was also evident when comparing the ketanserin + psilocybin group to the saline control group (Fig. S2h, p=0.431). We therefore conclude that the effects of psilocybin are at least partly mediated by its action at 5HT2A receptors.

**Figure 5.**
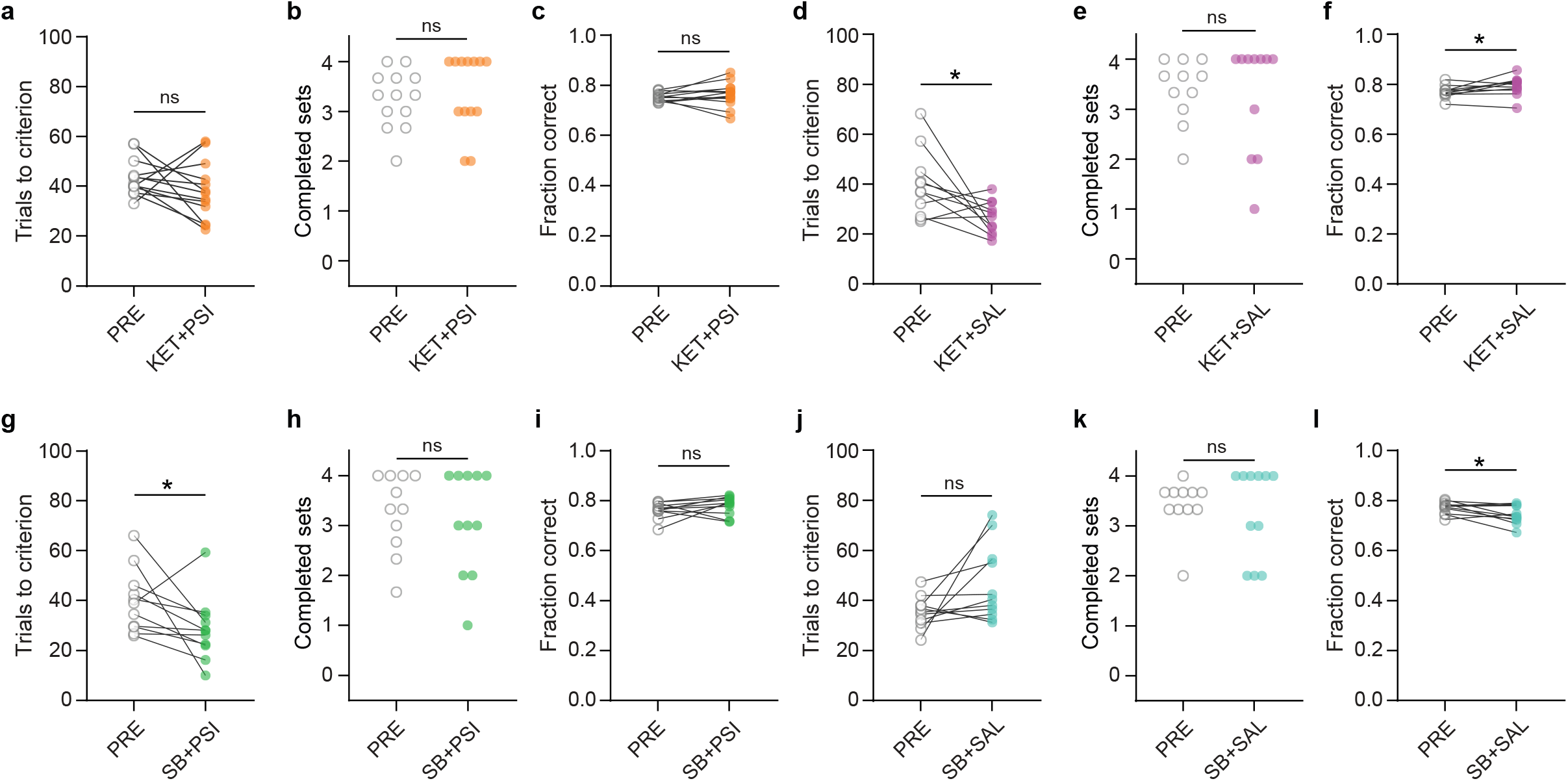
Pre-treatment with the 5HT2A antagonist ketanserin blocks psilocybin’s effect on cognitive flexibility. **(a)** Average trials-to-criterion for baseline (PRE, gray open circles) and ketanserin-psilocybin condition (KET+PSI, orange circles, n=14). **(b)** Number of completed sets in baseline and ketanserin-psilocybin conditions. **(c)** Fraction of correct responses averaged across all sets in baseline and ketanserin-psilocybin conditions. **(d)** Average trials-to-criterion for baseline (PRE, gray open circles) and ketanserin-saline conditions (KET+SAL, pink circles, n=11). * p=0.026, paired t-test. **(e)** Number of completed sets in baseline and ketanserin-saline conditions. **(f)** Fraction of correct responses averaged across all sets in baseline and ketanserin-saline conditions. **(g)** Average trials-to-criterion for baseline (PRE, gray open circles) and SB242084-psilocybin conditions (SB+PSI, green circles, n=11). **(h)** Number of completed sets in baseline and SB242084-psilocybin conditions. **(i)** Fraction of correct responses averaged across all sets in baseline and SB242084-psilocybin conditions. **(j)** Average trials-to-criterion for baseline (PRE, gray open circles) and SB242084-saline conditions (SB+SAL, teal circles, n=11). * p=0.032, Wilcoxon sign-rank test. **(k)** Number of completed sets in baseline and SB242084-saline conditions. **(l)** Fraction of correct responses averaged across all sets in baseline and SB242084- saline conditions. * p=0.031, paired t-test.

Ratsinjected with ketanserin followed by saline also showed a performance improvement comparable to psilocybin-treated animals. They required fewer trials to reach criterion (Fig. 5d, p=0.026, n=11), while completing a similar number of sets as during baseline (Fig. 5e, p=0.680), and showed a subtle but significant increase in the percent of correct trials (Fig. 5f, p=0.050). We also found a significant effect when comparing to the saline control group (Fig. S2h, p=0.0004). This effect of ketanserin is consistent with a previous report [34] and suggests a complex relationship between 5HT2A receptor function and cognitive flexibility.

Psilocin also binds to 5HT2C receptors. While ketanserin binds more strongly to 5HT2A receptors, it has moderate affinity at 5HT2C receptors. We, however, found that the 5HT2C antagonist SB242084 (1 mg/kg, i.p.) did not impact psilocybin’s enhancement of cognitive flexibility in the set-shifting task (Fig. 5g-I, n=11; trials to criterion, p=0.0322; completed sets, p=0.484; fraction of correct responses, p=0.249; see also Supplementary Fig. S5). Injection of SB242084 after saline produced a trend towards impaired performance (Fig 5j-l, n=11; trials to criterion, p=0.062; completed sets, p=0.762; fraction of correct responses, p=0.031). The same results were seen when comparing the SB242084 groups to the saline animals (Fig. S2h; SB242084 + saline, p=0.999; SB242084 + psilocybin, p=0.010). Thus, the improvement in cognitive flexibility seen with acute psilocybin treatment is likely independent of its action at 5HT2C receptors.

## Discussion

Psilocybin has received considerable attention for its therapeutic potential in treating anxiety and depression when combined with psychotherapy or other clinician-assisted interventions [1,5–7,35–37]. The biological mechanism underlying this effect is not well understood, but one interesting hypothesis is that psilocybin increases cognitive flexibility [1]. We investigated this idea and the pharmacological basis for psilocybin’s action using rats trained on an extradimensional set-shifting task. We find that psilocybin improves the ability to flexibly switch between learned action patterns. In contrast, psilocybin does not affect learning of new environmental contingencies. This effect of psilocybin did not generalize to the serotonergic hallucinogen DOI, which impaired cognitive flexibility, highlighting differences between psilocybin and other psychedelics. We further found that the 5HT2A/C antagonist ketanserin, but not the 5HT2C-selective antagonist SB242084, blocked the psilocybin-induced improvement in flexibility. These findings demonstrate a possible mechanism for the therapeutic action of psilocybin and provide a behavioral model for future investigation of the biological basis of the clinical efficacy of acute psilocybin or related novel compounds.

### Psilocybin selectively improves switching between known behavioral strategies

Cognitive flexibility in rodents has been studied using two experimental approaches: reversal learning tasks and set-shifting tasks. These two approaches, however, assess different constructs. Reversal learning requires animals to learn new stimulus-outcome or response-outcome associations, in Pavlovian and instrumental versions of the task respectively [33]. Set-shifting tasks, similar to the test we used here, instead involve an attentional and behavioral shift based on the representation of previously learned strategies [32]. While action-outcome associations do change in our task, the primary driver of behavior is the choice between internalized representationsof task rules and not newly learned associations. We note that it is possible that psilocybin would have different effects in an instrumental version of the reversal learning task, or one that does not include stimuli of both positive and negative valences, which may be closer to updating tasks used in humans. Nevertheless, our results support the idea that psilocybin’s acute effects differentially affect flexibly choosing between known strategies vs updating of environmental contingencies.

Our findings are consistent with human studies with ayahuasca [38] and psilocybin [19], and have interesting implications for understanding psilocybin’s effect on brain function. Set-shifting and reversal learning involve distinct neural circuits [39,40]. A large number of studies in humans, non-human primates and rodents have identified the orbitofrontal cortex (OFC) as the primary region required for reversal learning [33,41–43], while medial prefrontal cortex (mPFC), and particularly prelimbic cortex (PrL), is necessary for extradimensional set-shifting [44–48]. Neuronal dynamics in the PrL represent task rules, and these representations shift when animals adapt their behavior [29,30,49–51]. The anterior cingulate cortex (ACC), another subdivision of mPFC, has also been implicated in flexible behavior, though its specific role in the rat is likely different than that of the PrL [40,52–54]. In humans, functional connectivity of the ACC is affected by psilocybin, providing a potential link to changes in flexibility, though the mechanism remains unclear [19].

At the level of neuronal networks, psilocybin may impact set-shifting by modulating neuronal activity in mPFC both directly, via activation of 5HT receptors, as well as indirectly via its effects on neuromodulatory regions that project to frontal cortex. Recent results that show a preeminent role of inhibitory interneurons in mediating strategy switching [55,56] and increased expression of activity-dependent genes in inhibitory neurons in the mPFC following psilocybin administration [57] support this notion. Psilocybin also impacts the activity in the dorsal raphe nucleus (DRN) and the locus coeruleus (LC) [58–62]. These two inter-connected regions reciprocally regulate their firing, and provide serotonergic and noradrenergic inputs to cortical regions. These two neurotransmitters systems have been strongly implicated in flexible decision-making and set-shifting [63–66]. Studies with psilocin and related compounds such as lysergic acid diethylamide (LSD) suggest that psilocybin decreases DRN firing [58,60,67,68], which may play a role in modulating behavioral flexibility. Finally, systemic administration of psilocybin likely influences activity in other subcortical regions that are critical for motivated behavior, such as the ventral tegmental area (VTA)[69]. Future studies are required to understand how psilocybin and other psychedelics affect network dynamics within the DRN, LC and VTA, and in the broader context of their projections to OFC and mPFC circuits.

### Pharmacological site of action of psilocybin

We found that psilocybin-mediated increases in cognitive flexibility in rats require activation of 5HT2A receptors, but not of 5HT2C receptors. While there is a substantial literature on the impact of 5HT receptors on cognitive flexibility tasks, results are often contradictory [63]. Consistent with our experiments with 5HT2A and 5HT2C antagonists, ketanserin has been shown to improve strategy switching in a spatial extra-dimensional shifting task, while SB242084 did not affect performance in the same task [34]. In reversal learning tasks, results are mixed with 5HT2A antagonists or agonists impairing [23,26,70] or improving [71] performance, and 5HT2C antagonists having no effect or enhancing [72] performance.

Psilocybin and ketanserin both improved set-shifting performance, but this improvement was absent with concurrent administration of both drugs. There are several possible explanations for this effect. First, activating and blocking 5HT2A receptors may be leading to different downstream circuit changes that each result in improved flexibility. These would have to be competing circuits, given that concurrent administration of both drugs has no effect. Alternatively, the two opposing manipulations may be converging on the same output circuit. It is worth noting that psilocybin has moderate affinity for 5HT1A receptors, the activation of which is known to depress firing in the DRN, thus likely causing lower extracellular 5HT levels in downstream areas. Studies in 5HT-depleted rodent models however generally show either no effect [73,74] or impairments in flexible behavior [75–77], in contrast with our data. It is possible that psilocybin’s action at 5HT2A receptors in this context can compensate for the decrease in endogenous 5HT and increase behavioral flexibility. Conversely, ketanserin’s antagonistic effect at 5HT2A receptors may result in increased 5HT levels by disrupting inhibitory feedback control of DRN neurons [78]. Therefore, ketanserin may be beneficially impacting flexibility by increasing endogenous 5HT levels, akin to commonly used antidepressants, such as selective serotonin reuptake inhibitors (SSRIs), an effect which has also been shown to improve performance in reversal learning tasks [71,79]. Finally, ketanserin binds to other non-serotonergic receptors including adrenergic receptors [80], which may contribute to its positive effects on set-shifting when administered with saline or psilocybin. Similarly, effects of psilocybin on behavioral flexibility may be independent of the 5HT2A receptors as is the case with its anti-anhedonic effects [21].

### Differences between psilocybin and DOI

Our findings that DOI has an opposite effect compared to psilocybin highlight significant differences between these drugs and urge caution in generalizing results obtained with DOI to other serotonergic psychedelics. DOI has been used in several studies to investigate the effect of 5HT2A receptor activation and as a readily available proxy for psychedelic function [68,81,82]. However, while psilocybin improved flexibility in our set-shifting task, DOI severely impaired performance. It is likely that the effects of DOI on other moderately complex behaviors will differ from those of other psychedelics, despite the assumed similarity between these drugs. Given the complex pharmacological profiles of psychedelics, caution should be used when attempting to generalize results, even if two compounds have similar affinities for specific receptors. Interestingly, two recent studies reported impaired cognitive flexibility in human subjects that received LSD, in contrast with the data on psilocybin [19,83,84]. These differences may be due to the subtle pharmacological differences between these drugs: LSD, like DOI, has higher affinity for 5HT2A than 5HT2C, while psilocin has more similar affinity to both receptors; LSD and psilocin both have moderately high affinity at 5HT1A receptors, while DOI does not; and LSD also binds to dopamine D1 and D2 receptors [85–88]. The clinical success of psilocybin may then be due to its very specific pharmacological profile, and may not generalize to other psychedelic compounds.

### Increased cognitive flexibility as a therapeutic mechanism for psilocybin

Our data show that an acute dose of psilocybin enhances cognitive flexibility. How could this effect support psilocybin’s therapeutic action? Our hypothesis is that psilocybin transiently counters the cognitive biases commonly found in individuals suffering from psychiatric disorders, particularly MDD. One caveat of our study is that we used healthy rats, not a preclinical model of MDD, so the results should be interpreted cautiously. Nevertheless, cognitive flexibility is associated with effective emotional regulation and general psychological health in healthy adults [89,90], and cognitive rigidity is a hallmark of depression and its associated perseverative cognitive processes such as rumination, all-or-nothing thinking, and acceptance of maladaptive beliefs about the self and environment [14,17,91]. Impaired cognitive flexibility is reported in MDD [92–96] and it has a substantial impact on treatment outcome [97,98]. Specific deficits in set-shifting are known to be present in patients with MDD [92], and, in preclinical models, antidepressant treatment ameliorates set-shifting impairments [99,100]. There is also a reported link in humans between increased cognitive flexibility and reduction of depressive symptomatology [91].

Psilocybin’s acute effect on cognitive flexibility may therefore provide a path for countering depressive symptoms. While our data do not address whether psilocybin results in a long-lasting increase in cognitive flexibility, even a transient effect during clinician-assisted intervention could be beneficial given that enhanced cognitive flexibility is advantageous during psychotherapy [101]. Current clinical protocols for psilocybin therapy involve extensive professional support and clinical-guided intervention [9]. We propose that the cognitive state brought about by psilocybin is especially favorable to receive therapeutic benefits from these treatments. This would support the current ideas that the set, the setting and the content of psilocybin administration sessions are all essential to achieve therapeutic benefits: psilocybin, by enhancing cognitive flexibility, may simply allow for a more effective intervention.

## Supporting information

All supplementary materials

## Author contributions

RJO, ATP and BM conceptualized and designed experiments. RJO, GG and ATP performed experiments. ATP analyzed data. ATP and BM wrote the paper.

## References

1. Andersen KAA, Carhart-Harris R, Nutt DJ, Erritzoe D. Therapeutic effects of classic serotonergic psychedelics: A systematic review of modern-era clinical studies. Acta Psychiatr Scand. 2021;143:101– 118.

2. Davis AK, Barrett FS, May DG, Cosimano MP, Sepeda ND, Johnson MW, et al. Effects of Psilocybin-Assisted Therapy on Major Depressive Disorder: A Randomized Clinical Trial. JAMA Psychiatry. 2021;78:481.

3. De Gregorio D, Aguilar-Valles A, Preller KH, Heifets BD, Hibicke M, Mitchell J, et al. Hallucinogens in Mental Health: Preclinical and Clinical Studies on LSD, Psilocybin, MDMA, and Ketamine. J Neurosci. 2021;41:891–900.

4. dos Santos RG, Osório FL, Crippa JAS, Riba J, Zuardi AW, Hallak JEC. Antidepressive, anxiolytic, and antiaddictive effects of ayahuasca, psilocybin and lysergic acid diethylamide (LSD): a systematic review of clinical trials published in the last 25 years. Therapeutic Advances in Psychopharmacology. 2016;6:193–213.

5. Grob CS, Danforth AL, Chopra GS, Hagerty M, McKay CR, Halberstadt AL, et al. Pilot Study of Psilocybin Treatment for Anxiety in Patients With Advanced-Stage Cancer. Arch Gen Psychiatry. 2011;68:71.

6. Carhart-Harris RL, Bolstridge M, Day CMJ, Rucker J, Watts R, Erritzoe DE, et al. Psilocybin with psychological support for treatment-resistant depression: six-month follow-up. Psychopharmacology. 2018;235:399–408.

7. Daws RE, Timmermann C, Giribaldi B, Sexton JD, Wall MB, Erritzoe D, et al. Increased global integration in the brain after psilocybin therapy for depression. Nat Med. 2022;28:844–851.

8. Griffiths RR, Johnson MW, Carducci MA, Umbricht A, Richards WA, Richards BD, et al. Psilocybin produces substantial and sustained decreases in depression and anxiety in patients with life-threatening cancer: A randomized double-blind trial. J Psychopharmacol. 2016;30:1181–1197.

9. Johnson M, Richards W, Griffiths R. Human hallucinogen research: guidelines for safety. J Psychopharmacol. 2008;22:603–620.

10. Phelps J. Developing Guidelines and Competencies for the Training of Psychedelic Therapists. Journal of Humanistic Psychology. 2017;57:450–487.

11. Kwan AC, Olson DE, Preller KH, Roth BL. The neural basis of psychedelic action. Nat Neurosci. 2022;25:1407–1419.

12. Carhart-Harris R, Nutt D. Serotonin and brain function: a tale of two receptors. J Psychopharmacol. 2017;31:1091–1120.

13. Carhart-Harris RL, Friston KJ. REBUS and the Anarchic Brain: Toward a Unified Model of the Brain Action of Psychedelics. Pharmacol Rev. 2019;71:316–344.

14. Dajani DR, Uddin LQ. Demystifying cognitive flexibility: Implications for clinical and developmental neuroscience. Trends in Neurosciences. 2015;38:571–578.

15. Millan MJ, Agid Y, Brüne M, Bullmore ET, Carter CS, Clayton NS, et al. Cognitive dysfunction in psychiatric disorders: characteristics, causes and the quest for improved therapy. Nat Rev Drug Discov. 2012;11:141–168.

16. Snyder HR. Major depressive disorder is associated with broad impairments on neuropsychological measures of executive function: A meta-analysis and review. Psychological Bulletin. 2013;139:81–132.

17. Southwick SM, Vythilingam M, Charney DS. The Psychobiology of Depression and Resilience to Stress: Implications for Prevention and Treatment. Annu Rev Clin Psychol. 2005;1:255–291.

18. Stange JP, Alloy LB, Fresco DM. Inflexibility as a Vulnerability to Depression: A Systematic Qualitative Review. Clin Psychol Sci Pract. 2017;24:245–276.

19. Doss MK, Považan M, Rosenberg MD, Sepeda ND, Davis AK, Finan PH, et al. Psilocybin therapy increases cognitive and neural flexibility in patients with major depressive disorder. Transl Psychiatry. 2021;11:574.

20. Bale TL, Abel T, Akil H, Carlezon WA, Moghaddam B, Nestler EJ, et al. The critical importance of basic animal research for neuropsychiatric disorders. Neuropsychopharmacol. 2019;44:1349–1353.

21. Hesselgrave N, Troppoli TA, Wulff AB, Cole AB, Thompson SM. Harnessing psilocybin: antidepressant-like behavioral and synaptic actions of psilocybin are independent of 5-HT2R activation in mice. Proc Natl Acad Sci USA. 2021;118:e2022489118.

22. Shao L-X, Liao C, Gregg I, Davoudian PA, Savalia NK, Delagarza K, et al. Psilocybin induces rapid and persistent growth of dendritic spines in frontal cortex in vivo. Neuron. 2021;109:2535-2544.e4.

23. Amodeo DA, Hassan O, Klein L, Halberstadt AL, Powell SB. Acute serotonin 2A receptor activation impairs behavioral flexibility in mice. Behavioural Brain Research. 2020;395:112861.

24. Odland AU, Kristensen JL, Andreasen JT. The selective 5-HT2A receptor agonist 25CN-NBOH does not affect reversal learning in mice. Behavioural Pharmacology. 2021;32:448–452.

25. Darmani NA, Martin BR, Pandey U, Glennon RA. Do functional relationships exist between 5-HT1A and 5-HT2 receptors? Pharmacology Biochemistry and Behavior. 1990;36:901–906.

26. Boulougouris V, Glennon JC, Robbins TW. Dissociable Effects of Selective 5-HT2A and 5-HT2C Receptor Antagonists on Serial Spatial Reversal Learning in Rats. Neuropsychopharmacol. 2008;33:2007–2019.

27. Winstanley CA, Theobald DEH, Dalley JW, Glennon JC, Robbins TW. 5-HT2A and 5-HT2C receptor antagonists have opposing effects on a measure of impulsivity: interactions with global 5-HT depletion. Psychopharmacology. 2004;176:376–385.

28. Darrah JM, Stefani MR, Moghaddam B. Interaction of N-methyl-D-aspartate and group 5 metabotropic glutamate receptors on behavioral flexibility using a novel operant set-shift paradigm. Behavioural Pharmacology. 2008;19:225–234.

29. Del Arco A, Park J, Wood J, Kim Y, Moghaddam B. Adaptive Encoding of Outcome Prediction by Prefrontal Cortex Ensembles Supports Behavioral Flexibility. J Neurosci. 2017;37:8363–8373.

30. Park J, Wood J, Bondi C, Del Arco A, Moghaddam B. Anxiety Evokes Hypofrontality and Disrupts Rule-Relevant Encoding by Dorsomedial Prefrontal Cortex Neurons. Journal of Neuroscience. 2016;36:3322–3335.

31. Kim Y-B, Matthews M, Moghaddam B. Putative γ-aminobutyric acid neurons in the ventral tegmental area have a similar pattern of plasticity as dopamine neurons during appetitive and aversive learning: Associative learning in VTA. European Journal of Neuroscience. 2010;32:1564–1572.

32. Bissonette GB, Roesch MR. Neurophysiology of rule switching in the corticostriatal circuit. Neuroscience. 2017;345:64–76.

33. Izquierdo A, Brigman JL, Radke AK, Rudebeck PH, Holmes A. The neural basis of reversal learning: An updated perspective. Neuroscience. 2017;345:12–26.

34. Baker PM, Thompson JL, Sweeney JA, Ragozzino ME. Differential effects of 5-HT2A and 5-HT2C receptor blockade on strategy-switching. Behavioural Brain Research. 2011;219:123–131.

35. Davis AK, Barrett FS, May DG, Cosimano MP, Sepeda ND, Johnson MW, et al. Effects of Psilocybin-Assisted Therapy on Major Depressive Disorder: A Randomized Clinical Trial. JAMA Psychiatry. 2021;78:481.

36. dos Santos RG, Osório FL, Crippa JAS, Riba J, Zuardi AW, Hallak JEC. Antidepressive, anxiolytic, and antiaddictive effects of ayahuasca, psilocybin and lysergic acid diethylamide (LSD): a systematic review of clinical trials published in the last 25 years. Therapeutic Advances in Psychopharmacology. 2016;6:193–213.

37. Griffiths RR, Johnson MW, Carducci MA, Umbricht A, Richards WA, Richards BD, et al. Psilocybin produces substantial and sustained decreases in depression and anxiety in patients with life-threatening cancer: A randomized double-blind trial. J Psychopharmacol. 2016;30:1181–1197.

38. Murphy-Beiner A, Soar K. Ayahuasca’s ‘afterglow’: improved mindfulness and cognitive flexibility in ayahuasca drinkers. Psychopharmacology. 2020;237:1161–1169.

39. Hamilton DA, Brigman JL. Behavioral flexibility in rats and mice: contributions of distinct frontocortical regions: Flexibility and frontocortical function. Genes, Brain and Behavior. 2015;14:4–21.

40. Kesner RP, Churchwell JC. An analysis of rat prefrontal cortex in mediating executive function. Neurobiology of Learning and Memory. 2011;96:417–431.

41. Dias R, Robbins TW, Roberts AC. Dissociation in prefrontal cortex of affective and attentional shifts. Nature. 1996;380:69–72.

42. Hornak J, O’Doherty J, Bramham J, Rolls ET, Morris RG, Bullock PR, et al. Reward-related Reversal Learning after Surgical Excisions in Orbito-frontal or Dorsolateral Prefrontal Cortex in Humans. Journal of Cognitive Neuroscience. 2004;16:463–478.

43. Izquierdo A, Suda R, Murray E. Bilateral Orbital Prefrontal Cortex Lesions in Rhesus Monkeys Disrupt Choices Guided by Both Reward Value and Reward Contingency. Journal of Neuroscience. 2004;24:7540–7548.

44. Birrell JM, Brown VJ. Medial Frontal Cortex Mediates Perceptual Attentional Set Shifting in the Rat. J Neurosci. 2000;20:4320–4324.

45. Floresco SB, Block AE, Tse MTL. Inactivation of the medial prefrontal cortex of the rat impairs strategy set-shifting, but not reversal learning, using a novel, automated procedure. Behavioural Brain Research. 2008;190:85–96.

46. McAlonan K, Brown VJ. Orbital prefrontal cortex mediates reversal learning and not attentional set shifting in the rat. Behavioural Brain Research. 2003;146:97–103.

47. Ragozzino ME. The Contribution of the Medial Prefrontal Cortex, Orbitofrontal Cortex, and Dorsomedial Striatum to Behavioral Flexibility. Annals of the New York Academy of Sciences. 2007;1121:355–375.

48. Rich EL, Shapiro ML. Prelimbic/Infralimbic Inactivation Impairs Memory for Multiple Task Switches, But Not Flexible Selection of Familiar Tasks. Journal of Neuroscience. 2007;27:4747–4755.

49. Bissonette GB, Roesch MR. Neural correlates of rules and conflict in medial prefrontal cortex during decision and feedback epochs. Front Behav Neurosci. 2015;9.

50. Durstewitz D, Vittoz NM, Floresco SB, Seamans JK. Abrupt Transitions between Prefrontal Neural Ensemble States Accompany Behavioral Transitions during Rule Learning. Neuron. 2010;66:438–448.

51. Rich EL, Shapiro M. Rat Prefrontal Cortical Neurons Selectively Code Strategy Switches. Journal of Neuroscience. 2009;29:7208–7219.

52. Ragozzino ME, Wilcox C, Raso M, Kesner RP. Involvement of rodent prefrontal cortex subregions in strategy switching. Behavioral Neuroscience. 1999;113:32–41.

53. Bissonette GB, Powell EM, Roesch MR. Neural structures underlying set-shifting: Roles of medial prefrontal cortex and anterior cingulate cortex. Behavioural Brain Research. 2013;250:91–101.

54. Brockett AT, Tennyson SS, deBettencourt CA, Gaye F, Roesch MR. Anterior cingulate cortex is necessary for adaptation of action plans. Proc Natl Acad Sci USA. 2020;117:6196–6204.

55. Cho KKA, Hoch R, Lee AT, Patel T, Rubenstein JLR, Sohal VS. Gamma Rhythms Link Prefrontal Interneuron Dysfunction with Cognitive Inflexibility in Dlx5/6+/− Mice. Neuron. 2015;85:1332–1343.

56. Cho KKA, Shi J, Sohal VS. Long-range inhibition synchronizes and updates prefrontal task activity. BioRxiv. 2022. 1 April 2022. https://doi.org/10.1101/2022.03.31.486626.

57. Davoudian PA, Shao L-X, Kwan AC. Shared and distinct brain regions targeted for immediate early gene expression by ketamine and psilocybin. BioRxiv. 2022. 20 March 2022. https://doi.org/10.1101/2022.03.18.484437.

58. Aghajanian GJ, Haigler HJ. Hallucinogenic indoleamines: Preferential action upon presynaptic serotonin receptors. Psychopharmacology Communications. 1975;1:619–629.

59. Aghajanian GK. Mescaline and LSD facilitate the activation of locus coeruleus neurons by peripheral stimuli. Brain Research. 1980;186:492–498.

60. Aghajanian GK, Foote WE, Sheard MH. Action of psychotogenic drugs on single midbrain raphe neurons. The Journal of Pharmacology and Experimental Therapeutics. 1970;171:178–187.

61. Aghajanian GK, Foote WE, Sheard MH. Lysergic Acid Diethylamide: Sensitive Neuronal Units in the Midbrain Raphe. Science. 1968;161:706–708.

62. Rasmussen K, Aghajanian GK. Effect of hallucinogens on spontaneous and sensory-evoked locus coeruleus unit activity in the rat: reversal by selective 5-HT2antagonists. Brain Research. 1986;385:395– 400.

63. Alvarez BD, Morales CA, Amodeo DA. Impact of specific serotonin receptor modulation on behavioral flexibility. Pharmacology Biochemistry and Behavior. 2021;209:173243.

64. Grossman CD, Bari BA, Cohen JY. Serotonin neurons modulate learning rate through uncertainty. Current Biology. 2022;32:586-599.e7.

65. Kehagia AA, Murray GK, Robbins TW. Learning and cognitive flexibility: frontostriatal function and monoaminergic modulation. Current Opinion in Neurobiology. 2010;20:199–204.

66. Yu AJ, Dayan P. Uncertainty, Neuromodulation, and Attention. Neuron. 2005;46:681–692.

67. Aghajanian GK, Vandermaelen CP. Intracellular recordings from serotonergic dorsal raphe neurons: pacemaker potentials and the effects of LSD. Brain Research. 1982;238:463–469.

68. Vollenweider FX, Kometer M. The neurobiology of psychedelic drugs: implications for the treatment of mood disorders. Nat Rev Neurosci. 2010;11:642–651.

69. De Gregorio D, Posa L, Ochoa-Sanchez R, McLaughlin R, Maione S, Comai S, et al. The hallucinogen d -lysergic diethylamide (LSD) decreases dopamine firing activity through 5-HT 1A, D 2 and TAAR 1 receptors. Pharmacological Research. 2016;113:81–91.

70. Amodeo DA, Jones JH, Sweeney JA, Ragozzino ME. Risperidone and the 5-HT _2A_ Receptor Antagonist M100907 Improve Probabilistic Reversal Learning in BTBR T + tf/J Mice: 5HT _2A_ receptor blockade improves reversal learning in BTBR mice. Autism Res. 2014;7:555–567.

71. Furr A, Lapiz-Bluhm MD, Morilak DA. 5-HT2A receptors in the orbitofrontal cortex facilitate reversal learning and contribute to the beneficial cognitive effects of chronic citalopram treatment in rats. Int J Neuropsychopharm. 2012;15:1295–1305.

72. Boulougouris V, Robbins TW. Enhancement of Spatial Reversal Learning by 5-HT2C Receptor Antagonism Is Neuroanatomically Specific. Journal of Neuroscience. 2010;30:930–938.

73. Carlson KS, Whitney MS, Gadziola MA, Deneris ES, Wesson DW. Preservation of Essential Odor-Guided Behaviors and Odor-Based Reversal Learning after Targeting Adult Brain Serotonin Synthesis. Eneuro. 2016;3:ENEURO.0257-16.2016.

74. Clarke HF, Walker SC, Crofts HS, Robbins TW, Roberts AC. Prefrontal Serotonin Depletion Affects Reversal Learning But Not Attentional Set Shifting. Journal of Neuroscience. 2005;25:532–538.

75. Lapiz-Bluhm MDS, Soto-Piña AE, Hensler JG, Morilak DA. Chronic intermittent cold stress and serotonin depletion induce deficits of reversal learning in an attentional set-shifting test in rats. Psychopharmacology. 2009;202:329–341.

76. van der Plasse G, Feenstra MGP. Serial reversal learning and acute tryptophan depletion. Behavioural Brain Research. 2008;186:23–31.

77. Wallace A, Pehrson AL, Sánchez C, Morilak DA. Vortioxetine restores reversal learning impaired by 5-HT depletion or chronic intermittent cold stress in rats. Int J Neuropsychopharm. 2014;17:1695– 1706.

78. Liu R, Jolas T, Aghajanian G. Serotonin 5-HT2 receptors activate local GABA inhibitory inputs to serotonergic neurons of the dorsal raphe nucleus. Brain Research. 2000;873:34–45.

79. Brigman JL, Mathur P, Harvey-White J, Izquierdo A, Saksida LM, Bussey TJ, et al. Pharmacological or Genetic Inactivation of the Serotonin Transporter Improves Reversal Learning in Mice. Cerebral Cortex. 2010;20:1955–1963.

80. Casey AB, Cui M, Booth RG, Canal CE. “Selective” serotonin 5-HT2A receptor antagonists. Biochemical Pharmacology. 2022;200:115028.

81. Celada P, Puig MV, Díaz-Mataix L, Artigas F. The Hallucinogen DOI Reduces Low-Frequency Oscillations in Rat Prefrontal Cortex: Reversal by Antipsychotic Drugs. Biological Psychiatry. 2008;64:392–400.

82. Michaiel AM, Parker PRL, Niell CM. A Hallucinogenic Serotonin-2A Receptor Agonist Reduces Visual Response Gain and Alters Temporal Dynamics in Mouse V1. Cell Reports. 2019;26:3475-3483.e4.

83. Pokorny T, Duerler P, Seifritz E, Vollenweider FX, Preller KH. LSD acutely impairs working memory, executive functions, and cognitive flexibility, but not risk-based decision-making. Psychol Med. 2020;50:2255–2264.

84. Wießner I, Olivieri R, Falchi M, Palhano-Fontes F, Oliveira Maia L, Feilding A, et al. LSD, afterglow and hangover: Increased episodic memory and verbal fluency, decreased cognitive flexibility. European Neuropsychopharmacology. 2022;58:7–19.

85. Halberstadt AL, Koedood L, Powell SB, Geyer MA. Differential contributions of serotonin receptors to the behavioral effects of indoleamine hallucinogens in mice. J Psychopharmacol. 2011;25:1548–1561.

86. Nichols DE. Psychedelics. Pharmacol Rev. 2016;68:264–355.

87. Watts VJ, Mailman RB, Lawler CP, Neve KA, Nichols DE. LSD and structural analogs: pharmacological evaluation at D1 dopamine receptors. Psychopharmacology. 1995;118:401–409.

88. Giacomelli S, Palmery M, Romanelli L, Cheng CY, Silvestrini B. Lysergic acid diethylamide (LSD) is a partial agonist of D2 dopaminergic receptors and it potentiates dopamine-mediated prolactin secretion in lactotrophs in vitro. Life Sciences. 1998;63:215–222.

89. Kashdan TB, Rottenberg J. Psychological flexibility as a fundamental aspect of health. Clinical Psychology Review. 2010;30:865–878.

90. Murphy FC, Michael A, Sahakian BJ. Emotion modulates cognitive flexibility in patients with major depression. Psychol Med. 2012;42:1373–1382.

91. Dennis JP, Vander Wal JS. The Cognitive Flexibility Inventory: Instrument Development and Estimates of Reliability and Validity. Cogn Ther Res. 2010;34:241–253.

92. Snyder HR. Major depressive disorder is associated with broad impairments on neuropsychological measures of executive function: A meta-analysis and review. Psychological Bulletin. 2013;139:81–132.

93. Austin M-P, Mitchell P, Goodwin GM. Cognitive deficits in depression: Possible implications for functional neuropathology. Br J Psychiatry. 2001;178:200–206.

94. Harvey PO, Le Bastard G, Pochon JB, Levy R, Allilaire JF, Dubois B, et al. Executive functions and updating of the contents of working memory in unipolar depression. Journal of Psychiatric Research. 2004;38:567–576.

95. Lee RSC, Hermens DF, Porter MA, Redoblado-Hodge MA. A meta-analysis of cognitive deficits in first-episode Major Depressive Disorder. Journal of Affective Disorders. 2012;140:113–124.

96. Taylor Tavares JV, Clark L, Cannon DM, Erickson K, Drevets WC, Sahakian BJ. Distinct Profiles of Neurocognitive Function in Unmedicated Unipolar Depression and Bipolar II Depression. Biological Psychiatry. 2007;62:917–924.

97. Papakostas GI. Cognitive Symptoms in Patients With Major Depressive Disorder and Their Implications for Clinical Practice. J Clin Psychiatry. 2014;75:8–14.

98. Trivedi MH, Greer TL. Cognitive dysfunction in unipolar depression: Implications for treatment. Journal of Affective Disorders. 2014;152–154:19–27.

99. Nikiforuk A, Popik P. Long-lasting cognitive deficit induced by stress is alleviated by acute administration of antidepressants. Psychoneuroendocrinology. 2011;36:28–39.

100. Lapiz MDS, Bondi CO, Morilak DA. Chronic Treatment with Desipramine Improves Cognitive Performance of Rats in an Attentional Set-Shifting Test. Neuropsychopharmacol. 2007;32:1000–1010.

101. Johnco C, Wuthrich VM, Rapee RM. The influence of cognitive flexibility on treatment outcome and cognitive restructuring skill acquisition during cognitive behavioural treatment for anxiety and depression in older adults: Results of a pilot study. Behaviour Research and Therapy. 2014;57:55–64.

