## Supplementary material for "Acute psilocybin enhances cognitive flexibility in rats": All supplementary materials

**Supplementary Methods**

*Animal training and use*

Animals were allowed to acclimate for 5-7 days after arrival from CRL and then handled for 10 min each day for 7 days. On day 4 food restriction began (12-14g for males, 10-12g for females, *ad libitum* water), until animals reached 85%-90% of their free-feeding weight. Thereafter the amount of chow was adjusted such that animals gained approximately 2-5g per week. Once at the target weight, rats were habituated to the operant chamber (Coulbourn Instruments) for 20 min per day for two days, with sucrose pellets (45 mg, Bio-Serv, Frenchtown, NJ) delivered at random times (5-20 second intervals) in conjunction with illumination of the food delivery port to signal reward availability. Specific training on each behavioral task then began.

For the set-shifting task, three separate cohorts of rats were used (n=12 per cohort, 6 of each sex; one female animal from cohort 2 was excluded prior to any injection due to aggressive behavior). Each drug treatment group generally included animals from different cohorts. Animals in cohort 1 and 2 were used for psilocybin, and DOI experiments. Animals in cohorts 2 and 3 were used for experiments with ketanserin and SB242084. The saline group included animals from all cohort. Note that there were no differences in baseline performance across groups, as shown in Supplementary Fig. S2h.

*Set-shifting task structure*

For the set-shifting task, rats were tested in an operant chamber containing a food delivery trough on one side and two nose-poke ports on the opposite side (Fig. 1a). On each trial one of the ports was illuminated, with the illumination sequence pseudo-randomized such that the light cue was not presented on the same side in more than two consecutive trials. On “Light rule” trials the illuminated port was rewarded, regardless of its spatial position; on “Side rule” trials, the port on one side (“Left” or “Right”) was rewarded, regardless of whether it was illuminated. A correct response resulted in illumination of the food port, and delivery of a sucrose pellet. An incorrect response resulted in illumination of the food port but no pellet delivery. The inter-trial interval (ITI, 10s fixed duration) began when the rat’s head was detected in the food port. At the end of the ITI a new trial immediately began. When a criterion of 10 consecutive correct responses was achieved, the rule switched such that the Light rule alternated with Side rule and each side rule occurred once per session (e.g. “Light”, “Left Side”, “Light”, “Right Side”). The starting rule alternated on every session. The sessions lasted until criterion was reached on all four sets or after a total time of 45 minutes.

*Behavioral training and testing on set-shifting task*

Training on the set-shifting task began with 4 daily sessions in which rats only experienced one rule, alternating between Light and Side rules. The first two sessions were 60 min long with fixed 5 sec inter-trial intervals (ITI), then the ITI was increased to 10 sec and the total length increased to 90 min. Rats were then trained to switch between rules, initially with a 60 min session length. Once they were able to complete four sets within this time, session time was reduced to 45 min. Performance under treatment conditions was assessed using a session length of 45 min with 10 sec ITIs and only after three consecutive days of stable baseline performance. Behavior was considered stable when in the past 6 sessions animals reached the following criteria: 70% correct response rate; number of completed sets differing by 1 set at most; a difference of less than 10% in the total number of trials to criterion for the session (Fig. 1c; see also ref. 28). Additionally, we required animals to complete at least 2 sets during each baseline day and at least 20 sessions prior to treatment. On treatment days, behavioral testing was performed 20 min after injection of the treatment drug. Antagonists were injected 10 min before the treatment drugs.

Rats were able to learn this task over several sessions. The basic principles of the task were in fact learned quickly, as evidenced in the examples shown in Fig. 1b. In the early sessions, animals were sometimes able to complete two sets (i.e. perform one shift), though they generally made many incorrect responses (Fig. 1b, top). Later sessions tended to have more completed sets, fewer incorrect responses, and fewer trials as a result (Fig. 1b, bottom). The example shown also highlights that in later sessions performance stabilized and animals took fewer trials to adapt their responses to the new rule. Thus, they became proficient at switching between light- and side-rule strategies once those strategies had been learned. To characterize performance in the task we plotted three different behavioral metrics across all of the sessions completed by animals in each cohort: the fraction of correct responses (Supplementary Fig. S1, top); the average number of trials to criterion across all sets completed in that session (Supplementary Fig. S1, middle); and the number of completed sets (Supplementary Fig. S1, bottom). For the fraction of correct responses and completed sets, performance rapidly improved in the first 15-20 sessions, before stabilizing from there onwards. This learning was less evident in the trials-to-criterion metric. The trials to criterion metric was defined as the number of trials the rat completed before reaching the criterion of 10 correct responses on a given set, averaged across all sets, and excluding incomplete sets. Because incomplete sets are excluded, this metric may be artificially low when animals quickly complete one set but then fail to shift their behavior. This will result in a low average value for trials to criterion but cannot be taken as evidence of effective set-shifting. For this reason, both the number of trials to criterion and the number of completed sets were used to evaluate set-shifting performance in the subsequent drug studies.

*Reversal learning task structure*

The reversal learning task was a modified version of a previously characterized task [28]. Prior to the experiment, food restriction, handling and habituation were performed as described above and animals were assigned to the saline (n=6, 3 males and 3 females) or psilocybin (n=7, 3 males and 4 females) group. The task was conducted in operant boxes (Coulbourn Instruments) with a light, tone and feeder port. The conditioned stimuli (CSs)for the task consisted of a blinking light and a 2.9 kHz tone. Each CS was associated with an appetitive (sucrose pellet, 45 mg) or aversive (mild footshock, 180ms, 0.2 mA) outcome (unconditioned stimulus, US), and these associations were counter-balanced across animals. Animals performed ten task sessions during which 50 stimulus-outcome associations for each kind of CS-US pairing (appetitive or aversive) were presented. Each CS was presented for 10 sec and followed by delivery of the US, immediately after which a 20 sec ITI began. On the sixth session, CS-US contingencies were reversed such that the CS that previously signaled an appetitive outcome now predicted the aversive one, and vice-versa. Injections of psilocybin (1 mg/kg) or an equivalent volume of saline were given 20 min before starting the task on the reversal session.

### Supplementary Figure S1

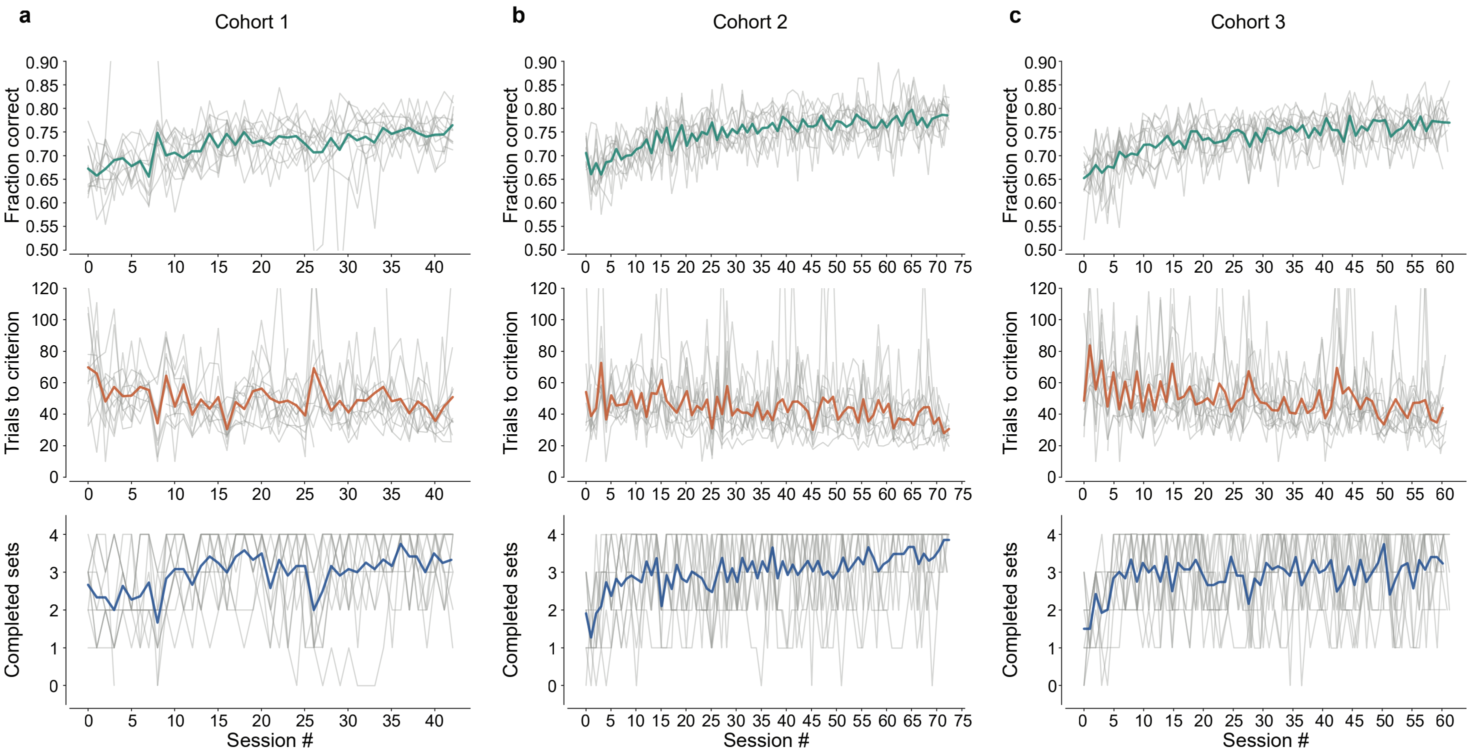

**Supplementary Figure S1.** Performance over time in the set-shifting task for each cohort. **(a)** Behavior in the set-shifting task over time, from the beginning of training until the last behavioral session, for a single cohort of animals (cohort 1, n=12, 6 males and 6 females). Top, fraction of correct trials across the whole session. Middle, average trials to criterion for the whole. Bottom, number of sets completed during each session. Gray lines show behavior of individual animals, thick colored lines show the average across animals. **(b)** As in (a), for cohort 2 (n=11, 6 males and 5 females). **(c)** As in (a), for cohort 3 (n=12, 6 males and 6 females).

### Supplementary Figure S2

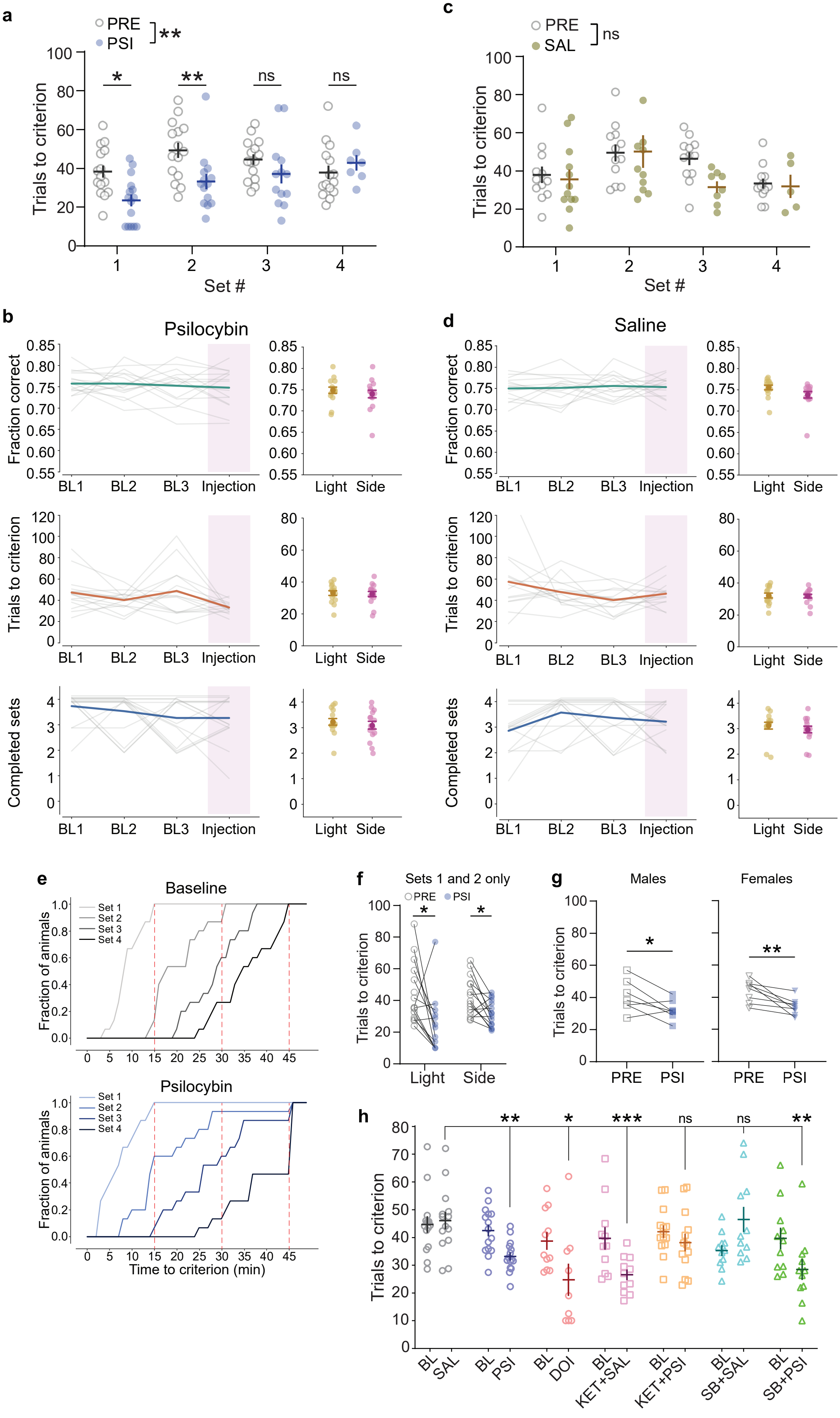

**Supplementary Figure S2.** Performance by set and stability for psilocybin condition. **(a)** Trials to criterion by set for baseline (PRE, gray) and psilocybin (PSI, blue) conditions. Circles represent data for individual animals, lines with error bars show mean ± SEM. Mixed-effects analysis: main effect of treatment, ** p = 0.006; interaction effect, Treatment x Session, p=0.029. Bonferroni post-hoc: * p = 0.010, ** p = 0.005. **(b)** Left: behavior for the three baseline sessions (BL1-3) and the drug injection session for animals receiving psilocybin as treatment, aligned to the injection session. Top, fraction of correct trials; middle, average number of trials to criterion across the session; bottom, number of complete sets (datapoints jittered for clarity). Gray lines indicate individual animals' behavior, thick colored lines denote average across animals. Right: Behavioral metrics for psilocybin-treated animals split by whether the session started with a Light (yellow) or Side (pink) rule, and averaged across all stable sessions (i.e. excluding the sessions during which animals are learning the task, see Figure 1c). Darker circle indicates mean ± SEM, lighter circles represent data for individual animals. **(c)** As in (a), for saline condition (brown circles). **(d)** As in (b), for saline condition. **(e)** Cumulative distribution across animals of time to criterion for each set in the baseline (top, grey lines) and psilocybin (bottom, blue lines) conditions. **(f)** Average trials to criterion for the first two sets in baseline (PRE, gray circles) and psilocybin (PSI, blue circles) conditions, split by rule (Light vs Side). * p<0.05, Wilcoxon sign-rank test with Bonferroni correction. **(g)** Average trials to criterion in baseline (PRE, gray) and psilocybin (PSI, blue) sessions, for male (n=7, left) and female (n=8, right) animals. Paired t-tests, * p=0.046, ** p=0.007. **(h)** Average trials to criterion for all treatment groups (SAL, grey circles; PSI, psilocybin, blue circles; DOI, red circles; KET+SAL, ketanserin and saline, pink squares; KET+PSI, ketanserin and psilocybin, orange squares; SB+SAL, SB242084 and saline, teal triangles; SB+PSI, SB242084 and psilocybin, green triangles; BL, baseline for each group). Two-way ANOVA, main effect of Timepoint (baseline or treatment) p=0.0019; main effect of Group, p=0.0002; interaction effect Timepoint x Group, p=0.0012. Post-hoc Dunnett's test comparing each group to saline control. No significant differences in the baseline timepoint. Results for treatment timepoints are shown on the graph: *p<0.05, ** p<0.01, ***p<0.001, ns p>0.4. Symbols represent data for individual animals, lines with error bars show mean ± SEM.

#### Supplementary Figure S3

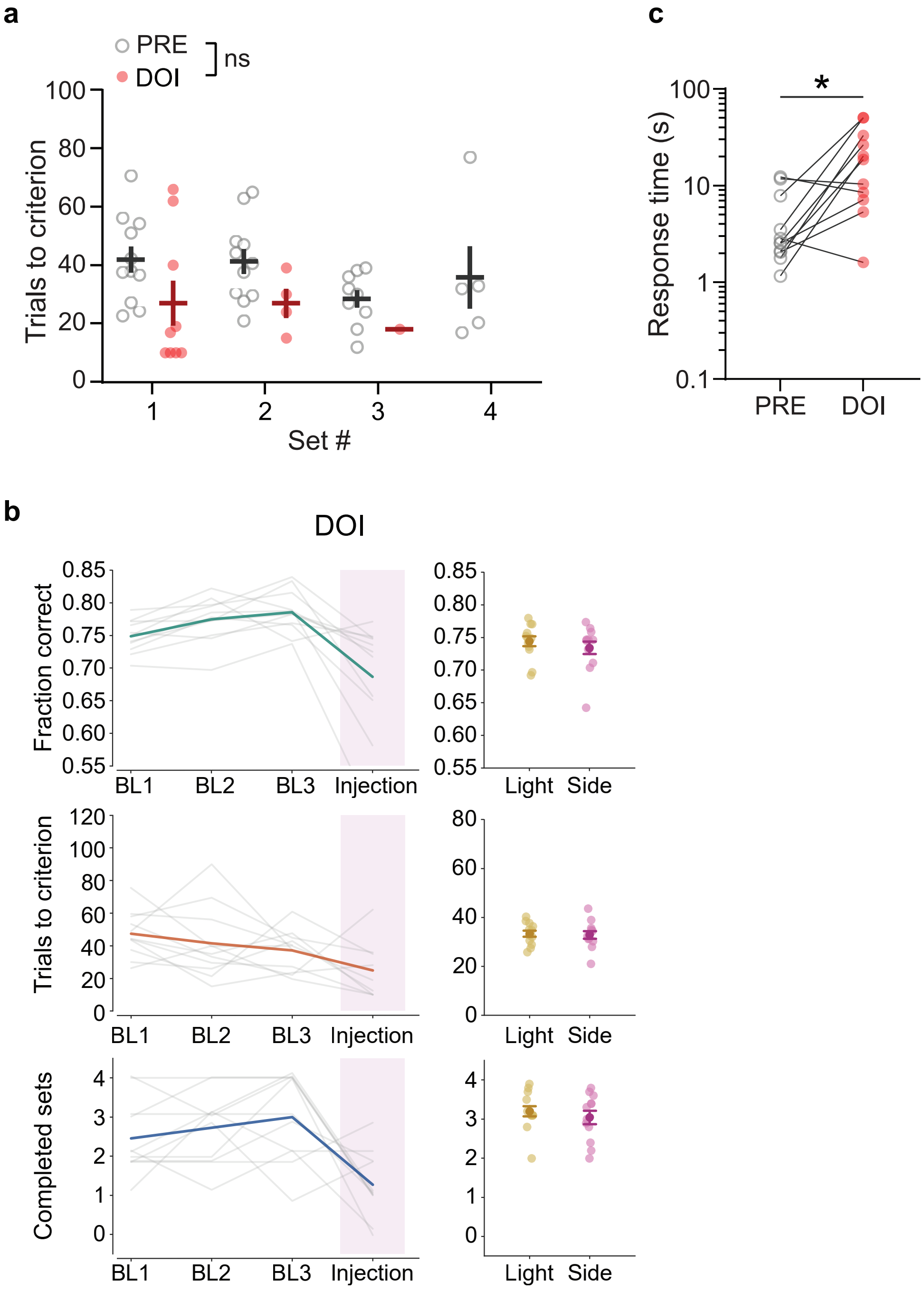

**Supplementary Figure S3.** Performance by set and stability for saline and DOI conditions. **(a)** Trials to criterion by set for baseline (PRE, gray open circles) and DOI (red circles). Circles represent data for individual animals, lines with error bars show mean ± SEM. **(b)** Left: Behavior for the three baseline sessions (BL1-3) and the drug injection session for animals receiving DOI as treatment, aligned to the injection session. Top, fraction of correct trials; middle, average number of trials to criterion across the session; bottom, number of complete sets (datapoints jittered for clarity). Gray lines indicate individual animals' behavior, thick colored lines denote average across animals. Right: Behavioral metrics for DOI-treated animals split by whether the session started with a Light (yellow) or Side (pink) rule, and averaged across all stable sessions. Darker circle indicates mean ± SEM, lighter circles represent data for individual animals. **(c)** Response time for baseline and DOI conditions. * p=0.019, Wilcoxon sign-rank test.

### Supplementary Figure S4

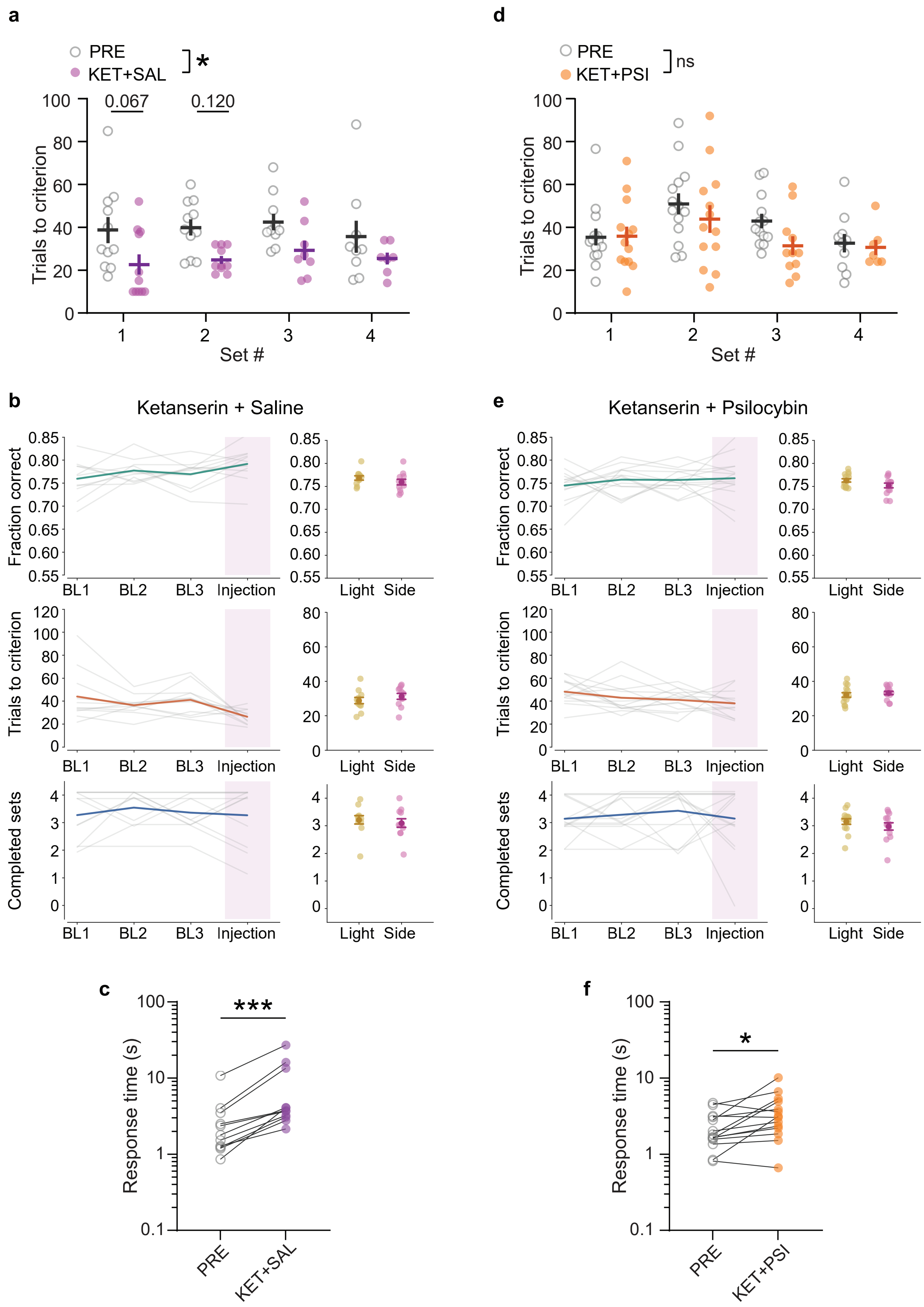

**Supplementary Figure S4.** Performance by set and stability for ketanserin-saline and ketanserin-psilocybin conditions. **(a)** Trials to criterion by set for baseline (PRE, gray open circles) and ketanserin-saline (KET+SAL, pink circles) conditions. Circles represent data for individual animals, lines with error bars show mean ± SEM. Mixed-effects analysis: main effect of treatment, * $p = 0.015$; Bonferroni post-hoc. **(b)** Left: Behavior for the three baseline sessions and the drug injection session for animals receiving ketanserin + saline as treatment, aligned to the injection session. Top, fraction of correct trials; middle, average number of trials to criterion across the session; bottom, number of complete sets (datapoints jittered for clarity). Gray lines indicate individual animals' behavior, thick colored lines denote average across animals. Right: Behavioral metrics for ketanserin-saline animals split by whether the session started with a Light (yellow) or Side (pink) rule, and averaged across all stable sessions. Darker circle indicates mean ± SEM, lighter circles represent data for individual animals. **(c)** Response time for baseline and ketanserin + saline conditions. *** $p=0.001$, Wilcoxon sign-rank test. **(d)** As in (a), for ketanserin-psilocybin condition (KET+PSI, orange circles). **(e)** As in (b), for ketanserin-psilocybin condition. **(f)** Response time for baseline and ketanserin + psilocybin conditions. * $p=0.011$, Wilcoxon sign-rank test.

#### Supplementary Figure S5

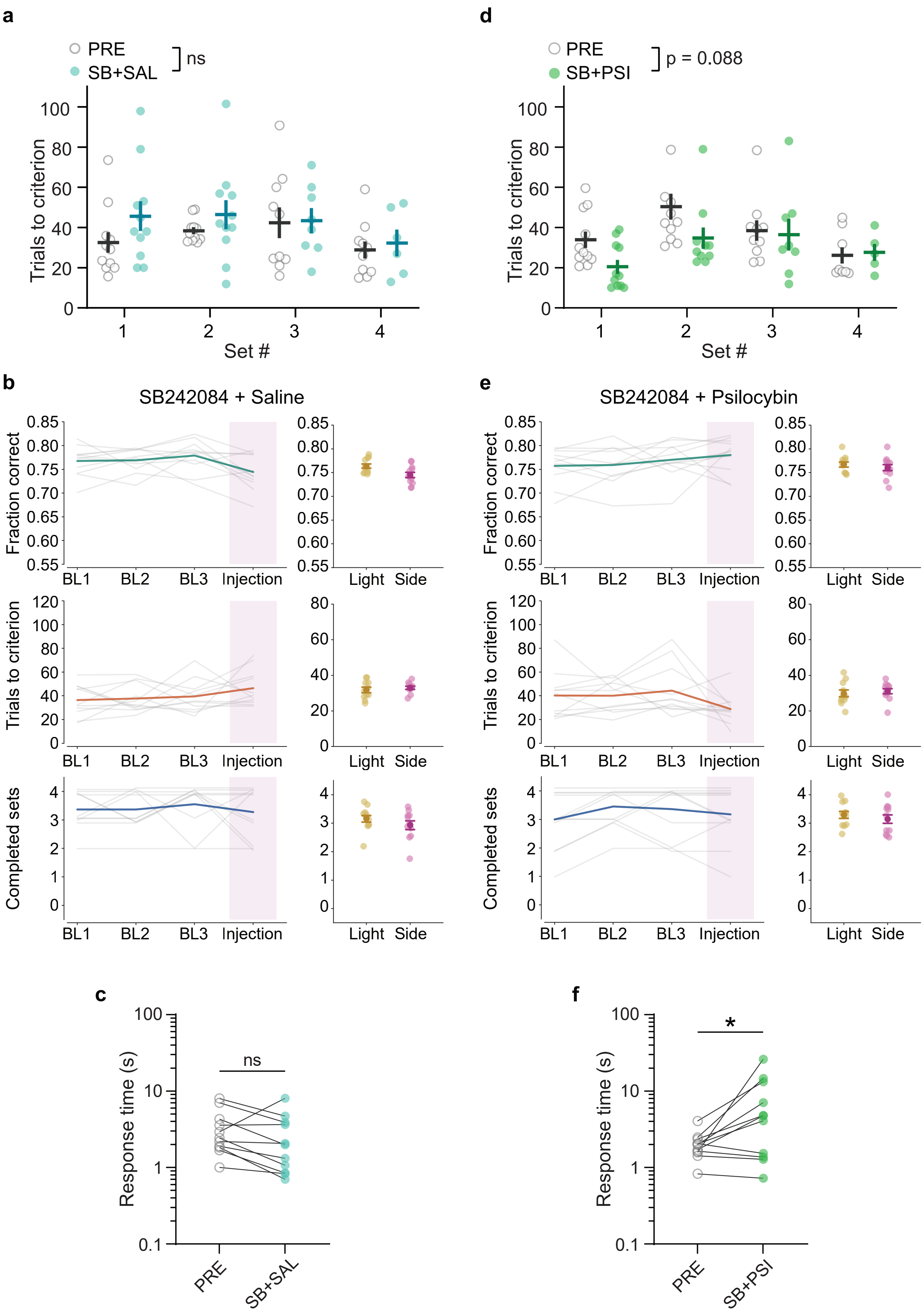

**Supplementary Figure S5.** Performance by set and stability for SB242084-saline and SB242084-psilocybin conditions. **(a)** Trials to criterion by set for baseline (PRE, gray open circles) and SB242084-saline (SB+SAL, teal circles) conditions. Circles represent data for individual animals, line with error bars show mean ± SEM. **(b)** Left: Behavior for the three baseline sessions and the drug injection session for animals receiving SB242084 + saline as treatment, aligned to the injection session. Top, fraction of correct trials; middle, average number of trials to criterion across the session; bottom, number of complete sets (datapoints jittered for clarity). Gray lines indicate individual animals' behavior, thick colored lines denote average across animals. Right: Behavioral metrics for psilocybin-treated animals split by whether the session started with a light (yellow) or side (pink) rule, and averaged across all stable sessions. Darker circle indicates mean ± SEM, lighter circles represent data for individual animals. **(c)** Response time for baseline and SB242084 + saline conditions. ns, $p=0.067$, Wilcoxon sign-rank test. **(d)** As in (a), for SB242084-psilocybin condition (SB+PSI, green circles). **(e)** As in (b), for SB242084-psilocybin condition. **(f)** Response time for baseline and SB242084 + psilocybin conditions. * $p=0.042$, Wilcoxon sign-rank test.

| ANIMAL # | 1ST INJECTION | 2ND INJECTION | 3RD INJECTION | 4TH INJECTION | FIGURES |
|---|---|---|---|---|---|
| **1** | PSI | DOI | | | 2, 4 |
| **2** | SAL | PSI | DOI | | 2, 4 |
| **3** | SAL | PSI | DOI | | 2, 4 |
| **4** | SAL | PSI | DOI | | 2, 4 |
| **5** | SAL | PSI | DOI | | 2, 4 |
| **6** | PSI | DOI | | | 2, 4 |
| **7** | SAL | PSI | DOI | | 2, 4 |
| **8** | SAL | PSI | DOI | | 2, 4 |
| **9** | SAL | PSI | DOI | | 2, 4 |
| **10** | SAL | PSI | DOI | | 2, 4 |
| **11** | PSI | DOI | | | 2, 4 |
| **12** | PSI | DOI | | | 2, 4 |
| **13** | PSI | KET+PSI | KET+SAL | SB+PSI | 2, 5 |
| **14** | KET+SAL | KET+PSI | | | 5 |
| **15** | KET+PSI | KET+SAL | SB+PSI | | 5 |
| **16** | SAL | KET+SAL | KET+PSI | | 2, 5 |
| **17** | KET+SAL | SB+PSI | | | 5 |
| **18** | PSI | KET+SAL | KET+PSI | | 2, 5 |
| **19** | KET+SAL | SB+PSI | | | 5 |
| **20** | SAL | KET+SAL | | | 2, 5 |
| **21** | PSI | KET+SAL | SB+PSI | | 2, 5 |
| **22** | SAL | KET+SAL | KET+PSI | | 2, 5 |
| **23** | SAL | KET+SAL | SB+PSI | | 2, 5 |
| **24** | KET+PSI | SB+PSI | SB+SAL | | 5 |
| **25** | SB+SAL | | | | 5 |
| **26** | KET+PSI | SB+SAL | | | 5 |
| **27** | KET+PSI | SB+PSI | SB+SAL | | 5 |
| **28** | KET+PSI | SB+SAL | | | 5 |
| **29** | SB+SAL | | | | 5 |
| **30** | KET+PSI | SB+SAL | | | 5 |
| **31** | KET+PSI | SB+PSI | SB+SAL | | 5 |
| **32** | KET+PSI | SB+PSI | | | 5 |
| **33** | SB+PSI | SB+SAL | | | 5 |
| **34** | SAL | KET+PSI | SB+SAL | | 2, 5 |
| **35** | SAL | SB+SAL | | | 2, 5 |

**Supplementary Table S1.** List of all animals used in the set-shifting experiments, along with the order of drugs received. The last column indicates to which figure's data that animal contributed to. SAL=saline; PSI=psilocybin; KET=ketanserin; SB=SB242084.
